# The PP2A phosphatase associates with the Arabidopsis TRAPPII tethering complex and dephosphorylates a TRAPPII-derived phosphopeptide *in vitro*

**DOI:** 10.64898/2026.09.03.749138

**Authors:** Alexander Strohmayr, Alexander Steiner, Christian Wiese, Miriam Abele, Eva Facher, Melina Altmann, Katia Belcram, Pascal Falter-Braun, Christina Ludwig, Martine Pastuglia, David Bouchez, Farhah F. Assaad

## Abstract

The transport protein particle II (TRAPPII) complex is a conserved regulator of post-Golgi membrane trafficking. In Arabidopsis, phosphorylation of the TRAPPII-specific subunit TRS120 by SHAGGY-like kinases modulates adaptive growth responses, but the phosphatases that reverse this phosphorylation remain unknown. Here, proteomic analyses identified subunits of Protein Phosphatase 2A (PP2A) in the TRAPPII interactome. PP2A subunits physically associated with TRAPPII, and double-mutant analyses revealed genetic interactions between PP2A and TRAPPII. Loss of TRAPPII function reduced the relative membrane association of PP2A scaffolding subunits. We established an *in vitro* assay for Arabidopsis PP2A holoenzyme activity using complexes transiently co-expressed and affinity-purified from *Nicotiana benthamiana*. Structural modelling and interface analysis predicted binding of a phosphorylated peptide encompassing a TRS120 phosphosite cluster at the PP2A catalytic interface, while biochemical assays showed that a PP2A holoenzyme containing the B2 regulatory subunit dephosphorylated this peptide. Together, these findings identify a B2-containing PP2A holoenzyme as a candidate phosphatase for TRS120 and support a model in which antagonistic SHAGGY-like kinase and PP2A activities couple signalling to membrane trafficking during plant development and environmental adaptation.

## INTRODUCTION

Plant growth requires the continuous coordination of cell division, cell expansion and cell differentiation, processes that notably depend on the precisely controlled delivery of proteins, lipids and cell-wall precursors to defined membrane domains (Delmer et al., 2024; Sablowski and Dornelas, 2014). At the trans-Golgi network/early endosome (TGN/EE), cargo sorting and membrane flow contribute to the establishment of membrane identity and to the regulation of endocytic and secretory pathways (Ravikumar et al., 2018). An important level of specificity is provided by multisubunit tethering complexes, which initiate the interaction between donor and recipient membranes prior to membrane fusion (Ravikumar et al., 2017). Among multisubunit tethering complexes, the transport protein particle II (TRAPPII) complex has emerged as a central regulator of post-Golgi trafficking in plants (Elliott et al., 2020; Kalde et al., 2019; Qi et al., 2011; Rybak et al., 2014).

Arabidopsis TRAPPII was first identified in forward and reverse genetic screens for cytokinesis-defective mutants: *trappii* mutants have a primary defect in cell plate biogenesis and feature multinucleate cells, cell-wall stubs, gaps and floating walls (Jaber et al., 2010; Thellmann et al., 2010). Subsequent work extended the role of TRAPPII to protein sorting, polarity, male gametophytic transmission, leaf-vein patterning, vascular development, and seedling development (Kalde et al., 2019; Naramoto et al., 2014; Qi et al., 2011; Rybak et al., 2014; Steiner et al., 2016; Wiese et al., 2024). Based on *Arabidopsis* interaction data and structural analogy with yeast TRAPPII, *Arabidopsis* TRAPPII is predicted to form a dimeric assembly of two equivalent multisubunit protomers, with an estimated molecular mass of approximately 1 MDa (Garcia et al., 2020; Kalde et al., 2019; Wiese et al., 2024). In this model, each protomer comprises an octameric shared core, together with the three TRAPPII-specific subunits AtTRS120, CLUB/AtTRS130, and the plant-specific subunit TRIPP (Garcia et al., 2020; Kalde et al., 2019). Indirect *in vivo* evidence suggests that Arabidopsis TRAPPII acts as a guanine-nucleotide exchange factor for Rab GTPases (Kalde et al., 2019), as has been shown in other species (Bagde and Fromme, 2022; Jenkins et al., 2020; Pinar et al., 2019).

Recent evidence indicates that Arabidopsis TRAPPII is subject to dynamic phosphorylation: the TRAPPII-specific subunit AtTRS120 is a substrate of multiple GSK3/SHAGGY-like kinases (AtSKs), including BRASSINOSTEROID INSENSITIVE 2 (BIN2), which collectively target three phosphosite clusters within plant-specific regions of AtTRS120 (Wiese et al., 2024). Functional analyses showed that TRS120 phosphorylation influences gravitropism, responses to osmotic stress and the mediation of root versus shoot growth trade-offs under combined stress (Kalbfuß et al., 2022; Wiese et al., 2024). Thus, while TRAPPII-kinase interactions have been characterised (Wiese et al., 2024), counteracting phosphatases have so far not been described. The emerging view of phosphorylation as a dynamic and reversible layer of plant signalling underscores the importance of identifying the phosphatases that establish phosphoprotein homeostasis by counterbalancing kinase activity (Praat et al., 2021; Vu et al., 2018). In this study, we identify a subset of protein phosphatases that interact with TRAPPII in Arabidopsis. We further establish a phosphatase-activity assay to test predicted phosphatase-TRAPPII enzyme-substrate relationships.

## RESULTS AND DISCUSSION

### PP2A subunits physically interact with TRAPPII

To identify phosphatase interactors of TRAPPII, we performed immunoprecipitation of Arabidopsis TRAPPII with untargeted mass spectrometry readout (IP-MS). Interestingly, numerous subunits of Protein Phosphatase 2A (PP2A) were identified in the TRAPPII interactome (Fig.1 A-B). PP2A is one of the major classes of serine/threonine phosphatases (Janssens and Goris, 2001) and has been implicated in BIN2/AtSK-regulated pathways (Kim et al., 2023; Tang et al., 2011). Across eukaryotes, PP2A holoenzymes have a conserved heterotrimeric organisation, comprising a catalytic C subunit, a scaffolding A subunit and a more diverse class of regulatory B subunits (Bheri and Pandey, 2019; Booker and DeLong, 2017). In *Arabidopsis thaliana*, the PP2A family is notably expanded, with the genome encoding 3 A (scaffold), 17 B (regulatory), and 5 C (catalytic) subunits (Fig. S1A). Of these, six were significantly enriched in IP-MS with the TRAPPII-specific subunit CLUB/AtTRS130 (Fig. 1A; Kalde et al., 2019). Because the TRAPPII-specific AtTRS120 subunit was identified as a substrate of the BIN2 SHAGGY-like kinase, IP-MS was performed with this subunit as well. We found that five PP2A subunits were significantly enriched in the TRS120 interactome (Fig. 1B). All three A isoforms were identified in the CLUB interactome (Fig. 1A), and all three were highly abundant in the TRS120 interactome (Fig. S1C). However, only A3 was significantly enriched in the AtTRS120 interactome (Fig. 1B). Arabidopsis PP2A regulatory B subunits are classified into three families, initially named after their molecular weights (Fig. S1A; Booker and DeLong, 2017; Máthé et al., 2023). The most significant enrichment obtained for PP2A B subunits in TRAPPII interactomes was for the close paralogs B1 and B2, the only members of the B55 subclade in Arabidopsis (Fig.1A-B; S1B, S1C). B regulatory subunits of the B56 and B72 subclades were found in TRAPPII interactomes, but at a lower abundance than B1 and B2 (Fig. S1B, S1C). For the five paralogs of the catalytic C subunit (C1-C5), C3 was found in the CLUB interactome, whereas C2 and C4 were identified in the AtTRS120 interactome (Fig. 1A-B). In both TRAPPII interactomes, the A and C subunits were the most enriched PP2A subunits (Fig. 1A-B), and A1, A2, A3 and C4 were as abundant as some TRAPPII subunits that co-purified with the TRS120 bait (Fig. S1C).

**Figure 1:**
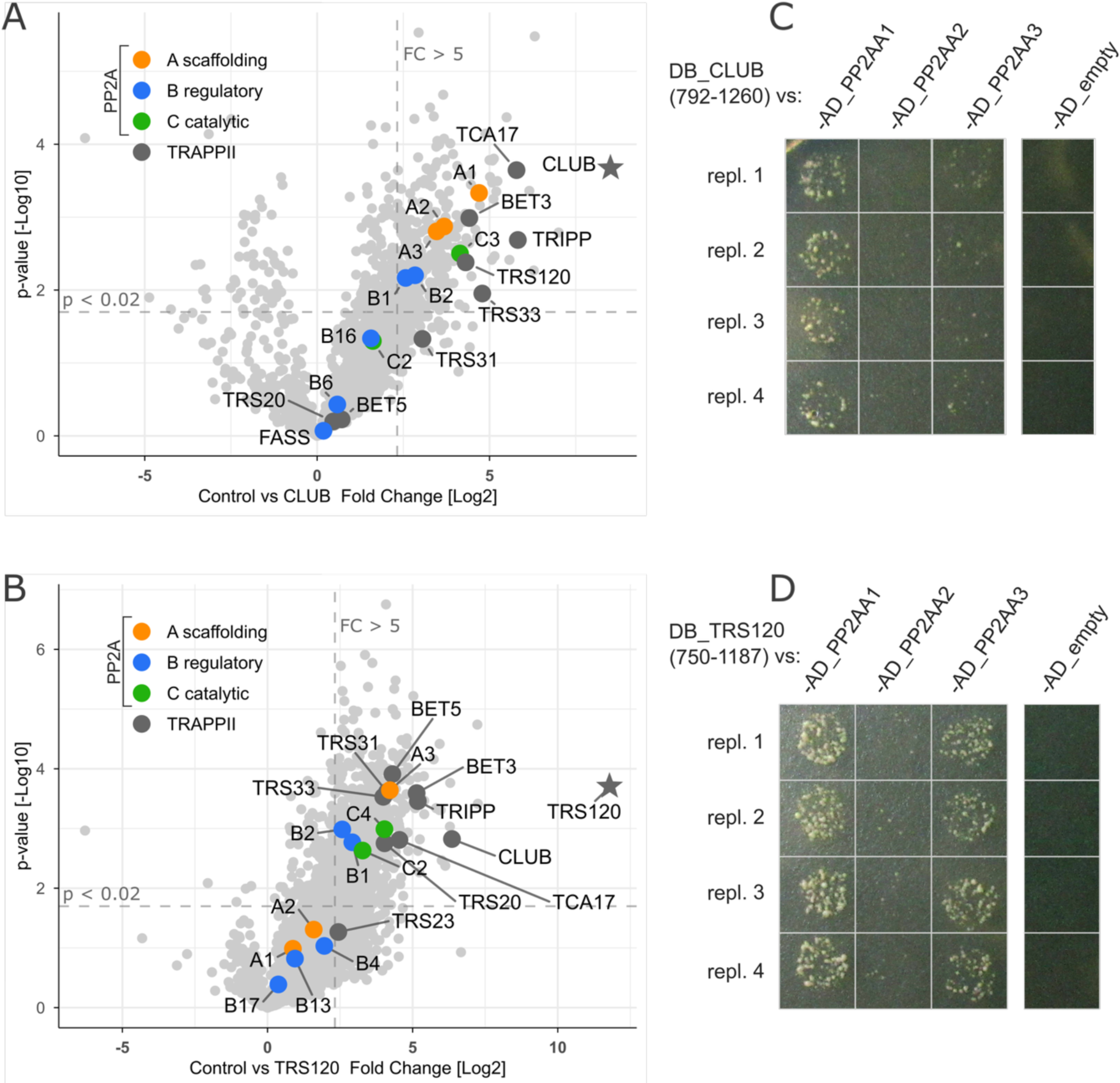
Physical interactions between TRAPPII and PP2A. **A, B.** IP-MS on inflorescences of TRAPPII-specific subunits CLUB/AtTRS130-GFP (A; new analysis of data previously described in Kalde *et al*., 2019) and AtTRS120-GFP (B; this study) searched against all Arabidopsis PP2A subunits (see Fig. S1A). The baits are denoted by stars. An empty vector cassette, with soluble GFP, was used as a negative control. Volcano plots are presented. The x-axis shows the fold change between the experiment and control (log2 scale), calculated as the difference between the mean label-free quantification (LFQ) values; positive values indicate enrichment with the TRAPPII bait. The y-axis shows the p-values of the LFQ signal in the experiment versus the control (negative log10 scale). See Fig. S1B, S1C for estimates of protein abundance. p-values were computed using a Welch two-sample t-test. Dotted grey lines represent cutoffs: p-value < 0.02 and fold change > 5 (log_2_(5) = 2.3). 6 PP2A subunits were significantly enriched in (A) and 5 in (B). **C, D.** Yeast two-hybrid assays using C-terminal fragments comprising the last 468 amino acids of CLUB/AtTRS130 (C) and the last 437 amino acids of AtTRS120 (D), with full-length open reading frames of all PP2AA scaffolding subunits in Arabidopsis thaliana. Positive interactions were scored between PP2AA1 and both TRAPPII truncations, and between PP2AA3 and AtTRS120.

Pairwise yeast two-hybrid experiments were performed between selected PP2A subunits (see Supplemental Information) and TRAPPII-specific truncations (Kalde et al., 2019; Rybak et al., 2014). This identified weak binary interactions between PP2AA1 and C-terminal fragments of CLUB/AtTRS130 and AtTRS120 (Fig. 1C, D). Furthermore, PP2AA3 interacted weakly with a C-terminal AtTRS120 fragment that encompasses the phosphosites targeted by the BIN2 kinase (Fig. 1D; Wiese et al., 2024). We conclude that the TRAPPII and PP2A complexes interact physically, with binary interactions between TRAPPII-specific and PP2AA scaffolding subunits.

### PP2A subunits interact genetically with TRAPPII

The TRAPPII tethering complex (Kalde et al., 2019; Thellmann et al., 2010; Wiese et al., 2024) and various members of the PP2A trimer (Camilleri et al., 2002; DeLong, 2006; Ren et al., 2022; Schaefer et al., 2017; Spinner et al., 2013; Yue et al., 2016) have been shown to be involved in cell division, cell expansion and development. To explore possible genetic interactions between TRAPPII and PP2A, and based on the Y2H experiments (Fig. 1C, D), we performed double-mutant analysis between TRAPPII (*trs120-4 and club-2)* and PP2AA3 (*pp2aa3-1* and *pp2aa3-2)* mutants.

We observed a partial suppression of the strong seedling-lethal phenotype of *trs120-4* in the double mutant with *pp2aa3-1,* with the double mutant having significantly longer roots (Fig. 2A, 2I, cf. Fig. 2E). The width and length of hypocotyl cells were assessed in scanning electron micrographs (SEMs) of single versus double mutants. Hypocotyl cell widths in the *trs120-4 pp2aa3-1* double mutant were significantly wider than in *trs120-4* single mutants, but did not significantly differ from *pp2aa3-1* single mutants (Fig. S2A). Cells were also significantly longer in the *trs120-4 pp2aa3-1* double mutant compared to *trs120-4,* but similar to *pp2aa3-1* (Fig. S2B). In contrast, we observed a significant reduction in hypocotyl cell length in *club-2 pp2aa3-2* in comparison to the single mutants (Fig. S2B). In fact, *club-2 pp2aa3-2* cells were almost isotropic (Fig. 2C, 2H, 2J; cf. Fig. 2B, 2G). We conclude that TRAPPII and PP2A interact genetically, with a synergistic PP2AA-CLUB/AtTRS130 interaction versus partial suppression with the AtTRS120 TRAPPII-specific subunit. Furthermore, *pp2a trappii* double mutants had defects in cell elongation and anisotropy.

**Figure 2.**
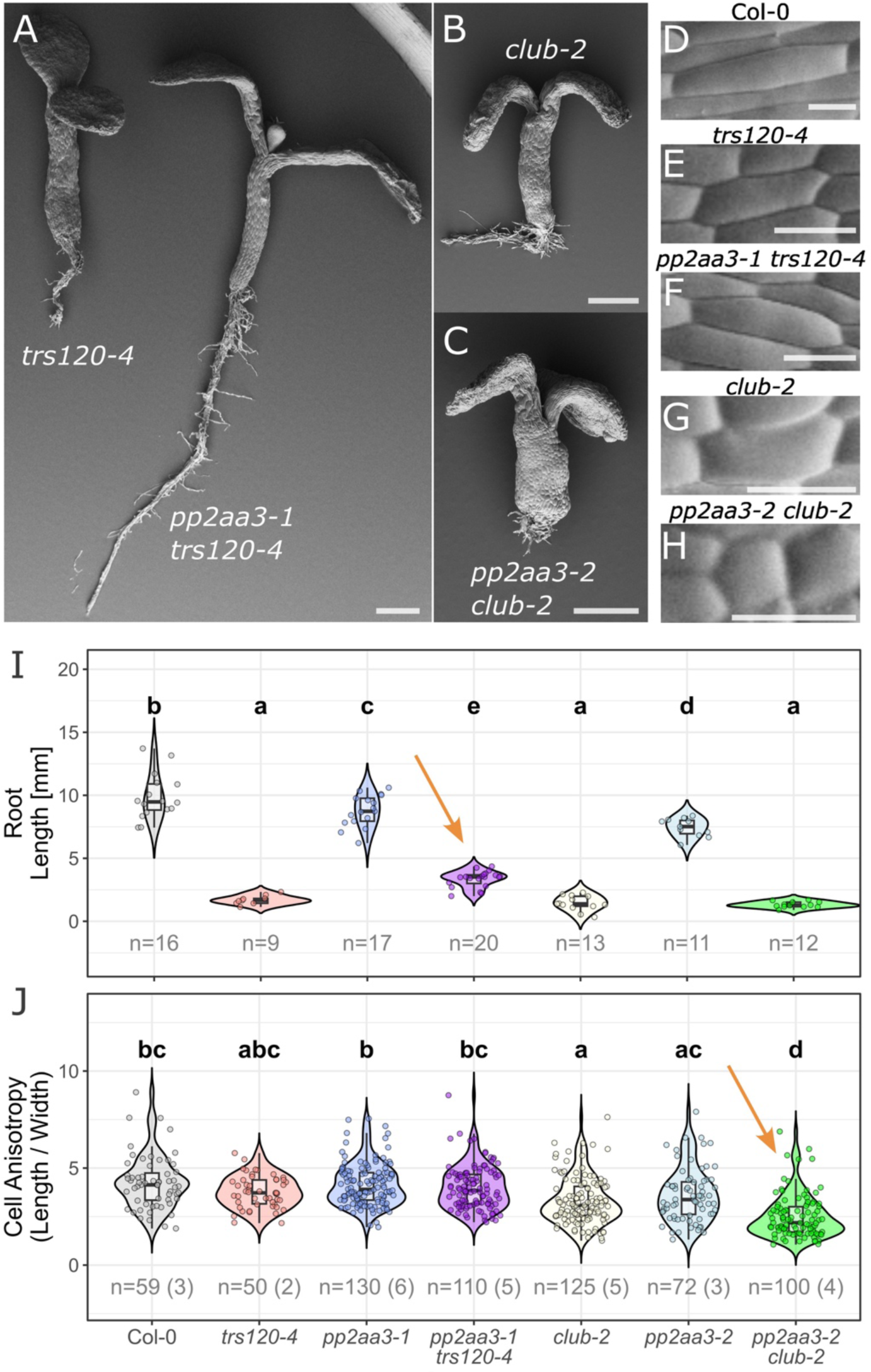
Genetic interactions between *trappii* and *pp2aa3* null mutants. **A-H**. Representative scanning electron micrographs of seedlings (A-C) or of hypocotyl cells (D-H): *trs120-4* (A, left; E); *pp2aa3-1 trs120-4* double mutant (A, right; F); *club-2* (B; G); *pp2aa3-2 club-2* double mutant (C; H); Col-0 (D). Scale bars: 500 µm for seedlings and 50 µm for close-ups. **I, J.** Violin–boxplot hybrids with overlaid single-cell or single-seedling measurements (jittered points) quantifying root length in mm (I) or cell anisotropy (length/width; J). Note the partial suppression of the *trs120-4* phenotype (A, left; I, red) in the *pp2aa3-1 trs120-4* double mutant (A, right; I, purple). *pp2aa3-1 trs120-4* had significantly longer cells (F; for quantification see Fig. S2B) and longer roots (I, orange arrow) than *trs120-4* (E; I, red). In contrast, note the enhancement of the *club-2* phenotype (B, G) in some *pp2aa3-2 club-2* double mutants (C); notably, the double mutant had significantly shorter (H; for quantification see Fig. S2B, green) and almost completely isodiametric cells, with values close to 1.0 (H; J, green; orange arrow). p-values were computed using either one-way ANOVA with Tukey’s HSD (I) or Kruskal–Wallis followed by Bonferroni-corrected Dunn tests (J). p-values are represented by compact letter displays. Sample sizes (n), shown in grey, indicate the number of cells (number of seedlings).

### Cortical microtubule alignment is impaired in *trappii* mutants

The cell morphology defects we observed in *pp2a trappii* double mutants are strongly suggestive of an impairment in microtubule-driven processes (Baskin, 2001; Spinner et al., 2013; Wasteneys and Ambrose, 2009). Indeed, cell morphology is at least partially controlled by interphase cortical microtubules that align with maximal tensile stress (Melogno et al., 2024). Given the central role of cortical microtubule organisation in establishing and maintaining cellular architecture, understanding how microtubule dynamics are integrated with membrane trafficking and signalling remains an important unresolved question (Müller, 2019) PP2A has been implicated in the organisation of cortical microtubule arrays during cell division and during interphase (Ren et al., 2022; Spinner et al., 2013). A GO term enrichment analysis of the TRAPPII interactome revealed a significant enrichment of proteins implicated in cytoskeletal organisation (Fig. S3). Therefore, we next investigated the microtubule cytoskeleton in *trappii* mutants. While *trappii* mutants have previously been shown to be impaired in phragmoplast microtubule array morphology (Steiner et al., 2016), interphase cortical arrays have not been studied in *trappii* alleles to date. Quantitative analysis of cortical microtubule co-alignment (Boudaoud et al., 2014) in *trappii* revealed that both *club-2 and trs120-4* epidermal root tip cells displayed a significantly lower cortical microtubule anisotropy than the wild type (Fig. 3A-B).

**Figure 3:**
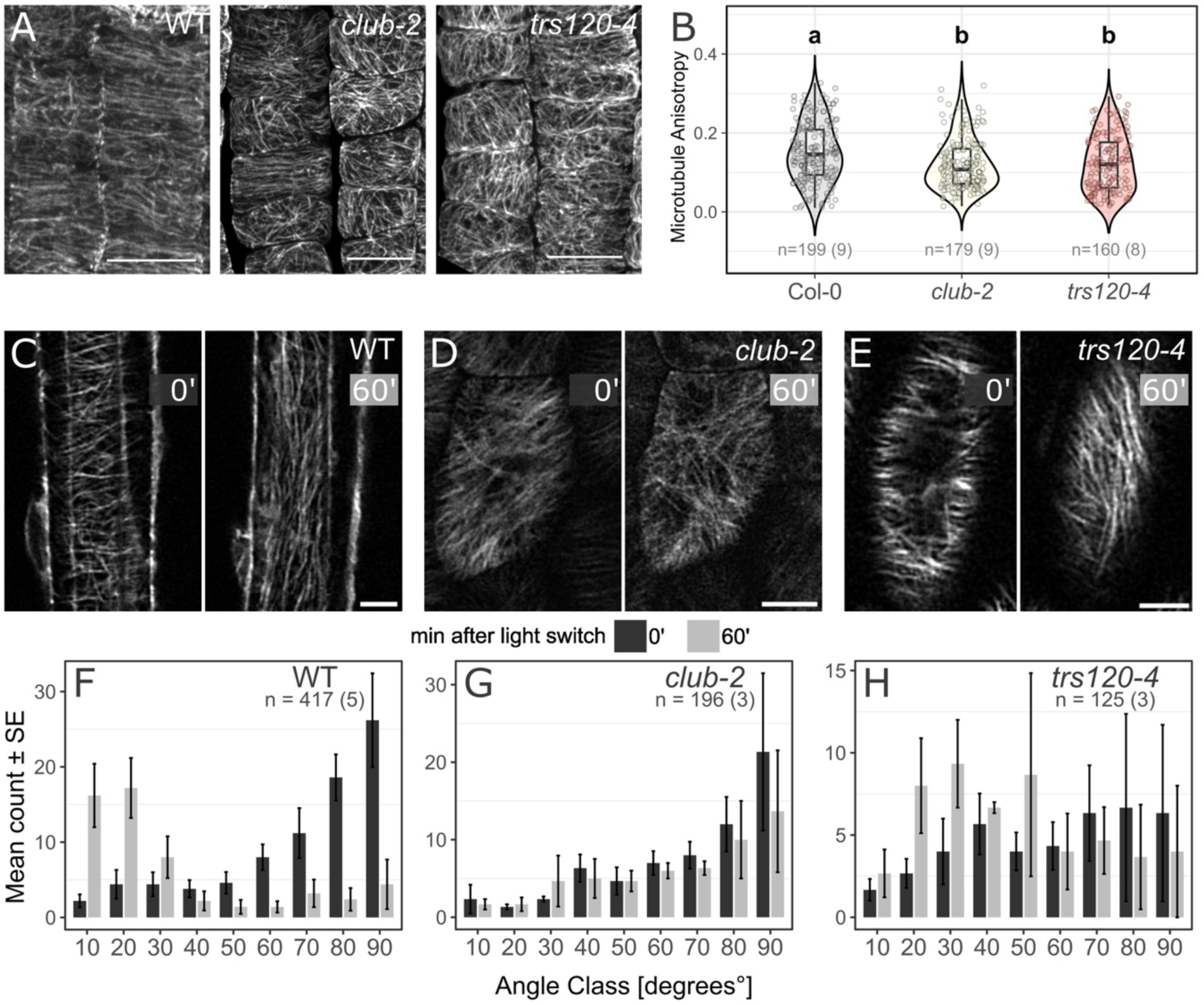
Cortical microtubules in *trappii* mutants. **A.** Immunolocalisation of tubulin in root tip epidermal cells. Images are maximum projections of cortical microtubules from the external cell faces (Col-0 (left), *club-2* (middle), *trs120-4* (right)). Scale bars: 10 µm. **B.** Semi-automated quantification of cortical microtubule (MT) ordering using FibrilTool (Boudaoud *et al*., 2014) in Col-0 wild type (grey), *club-2* (yellow), and *trs120-4* (red). Microtubule array anisotropy (the extent to which cortical microtubules displayed a dominant orientation, reflecting their degree of alignment, independently of the absolute orientation angle) was lower in the *club-2* and *trs120-4* mutants. Significance letters are derived from Holm-corrected pairwise Wilcoxon tests and planned two-group Wilcoxon comparisons. n = number of cells (number of root tips). **C-H.** Dynamic rearrangement of cortical microtubules in hypocotyl cells in *trappii* mutants **C, D, E.** Representative confocal images of the microtubule marker TUA5-mCherry before (left) and 60 min after the light stimulus (right). The same cell is depicted on the left (0’ for 0 min) and the right (60’ for 60 min). Scale bars: 10 µm. **F, G, H.** Quantification of microtubule orientation (0° for longitudinal and 90° for transverse), with classification in 10° angle bins and a computation of the number of MTs in each bin. Bars represent the mean ± SE (standard error of the mean) for the number of microtubules classified over different seedlings. Dark grey bars represent 0 min samples, and light grey bars represent the same sample 60 min after the light switch stimulus. n = number of cells (number of seedlings). **C, F.** Col-0. Note the homogeneous transverse to longitudinal realignment of cortical microtubules. **D, G.** *club-2.* The MTs were not responsive to the light stimulus and remained largely transverse. **E, H.** *trs120-4* displayed a more erratic behaviour of cortical microtubules before and after the light switch (see large error bars) and showed an incomplete reorientation.

The anomalies in *trappii* microtubule organisation prompted us to investigate whether the dynamic reorganisation of cortical microtubule arrays in response to an environmental stimulus was also affected. To this end, we used a TUA5-mCherry marker (Gutierrez et al., 2009) to observe microtubule realignment in response to a light stimulus in the hypocotyl cells of dark-grown seedlings (Sampathkumar et al., 2011; Ueda and Matsuyama, 2000). In dark-grown Arabidopsis hypocotyls, cortical microtubules organise into a transverse coaligned pattern that is critical for axial cell growth (Lucas et al., 2011; Vineyard et al., 2013). The switch from darkness to light causes a 90° realignment of the microtubules along the longitudinal axis, from transverse arrays in the dark to longitudinal arrays 60 min after the light stimulus (wild type; Fig. 3C, F). The orientation shift apparent in wild type was not observed in *club-2* and was incomplete and more variable in *trs120-4* (Fig. 3C–H). These observations indicate that static cortical array organisation and light-induced reorientation are impaired in *trappii* mutants. Similar cortical microtubule defects have been reported for *fass* (Kirik et al., 2012). As *FASS* encodes a B72-family regulatory subunit of PP2A (Fig. S1A), this finding is consistent with functional convergence between TRAPPII and PP2A.

### PP2A scaffolding subunits exhibit reduced membrane localisation in *trappii* mutants

Considering the cortical microtubule phenotypes observed in *trappii* and *pp2a* mutants (Ren et al., 2022; Schaefer et al., 2017; Spinner et al., 2013), we turned our attention to the cellular distribution of PP2A at the cortex of root meristem cells in *trappii* mutants. Previous work has shown that the Arabidopsis PP2A scaffolding subunits PP2AA1/RCN1 and PP2AA3 are not restricted to the cytosol but are also associated with cellular membranes (Blakeslee et al., 2008; Michniewicz et al., 2007). As these proteins lack intrinsic membrane-targeting sequences, their membrane localisation likely depends on interactions with membrane lipids or membrane-associated protein complexes (Blakeslee et al., 2008). We therefore tested the localisation of PP2AA1 and PP2AA3, the two PP2A A subunits interacting with TRS120 in yeast two-hybrid assays (Fig. 1D), in *trs120-4* mutant cells. We found that the overall fluorescence levels of YFP-PP2AA1 and YFP-PP2AA3 (Blakeslee et al., 2008) were reduced in *trs120-4* root meristem cells (Fig. 4A, 4C). Furthermore, the membrane localisation of PP2AA1 and PP2AA3 was affected (Fig. 4A, 4C), as evidenced by the reduced relative intensity of membrane versus cytosolic signal in *trs120-4* (Fig. 4B, 4D). We conclude that a functional TRAPPII complex contributes to the membrane association of PP2AA subunits, which may impact PP2A’s role in the organisation of cortical microtubule arrays.

**Figure 4.**
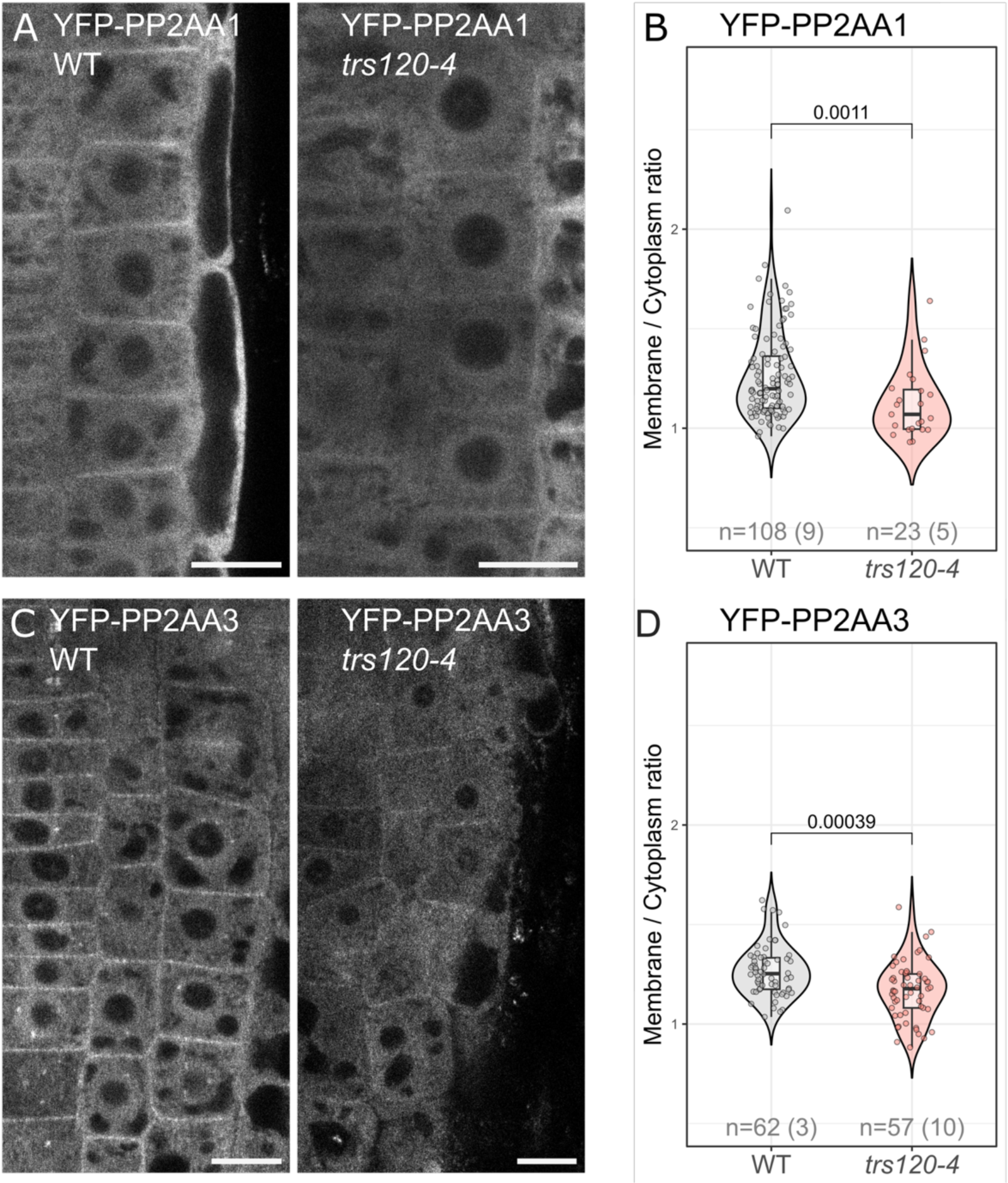
Localisation of PP2AA scaffolding subunits in *trs120-4*. Representative images **(A, C)** and violin plots quantifying the ratio of membrane to cytosolic signal intensity **(B, D)**. WT (left, grey in B, D) denotes phenotypically wild-type and *trs120-4* seedling-lethal (right, red in B, D) segregating seedlings in a hemizygous *trs120-4* line. Scale bars: 10 µm. **A, B.** ProPP2AA1:YFP-PP2AA1 (Blakeslee et al., 2008). **C, D.** ProPP2AA3:YFP-PP2AA3 (Blakeslee et al., 2008). The intensity values for membrane and cytoplasm were acquired by drawing equal rectangles in the respective areas. Only cells from the epidermis of the root meristem were considered for the quantification. Note the significant reduction of PP2A membrane localisation in *trs120-4*. p-values were computed with a Wilcoxon rank-sum test. n = number of cells (number of seedlings).

### A B2-containing PP2A complex dephosphorylates a TRS120-derived phosphopeptide

The physical and genetic interactions between TRAPPII and PP2A (Fig. 1, Fig. 2), and the altered distribution of PP2A in *trs120-4* (Fig. 4), prompted us to explore whether AtTRS120 is a direct PP2A substrate. The TRAPPII-specific AtTRS120 subunit has been shown to be differentially phosphorylated by SHAGGY-like kinases at three phosphosite clusters, referred to as α, β, and γ (Wiese et al., 2024; Table S2). To test whether TRS120 phosphosite status influences PP2A association, we compared PP2A recovery with phosphomimetic TRS120 variants. To this end, we mimicked phosphorylation by introducing serine-to-aspartate substitutions (S-to-D) at the α, β, and γ phosphosite clusters; the substitutions were introduced into genomic constructs by site-directed mutagenesis, and the resulting constructs were transformed into plants to generate stable GFP-tagged phosphovariant lines (Wiese et al., 2024). Co-immunoprecipitation of AtTRS120 phosphovariants with mass spectrometry readout showed that, of all 25 PP2A subunits surveyed, the regulatory B2 subunit was the only significantly enriched PP2A subunit in the AtTRS120 phosphomimetic-variant IP-MS datasets (Fig. 5A; Fig. S1A). Furthermore, this enrichment was observed solely with phosphomimetic substitutions at the TRS120 β-cluster (Fig. 5A). Together with the predominance of B1 and B2 in TRAPPII interactomes (Fig. 1A, 1B; Fig. S1B, S1C), these results suggest cluster-dependent differences in the association of B2 with TRS120 phosphovariants. Previous work implicates B2 (also referred to as B55β) in coordinating plant development with environmental signalling: it represses flowering, supports embryo viability, and is activated in response to ethylene to regulate root growth (Heidari et al., 2013; Heidari et al. 2023; Shao et al., 2022).

**Figure 5:**
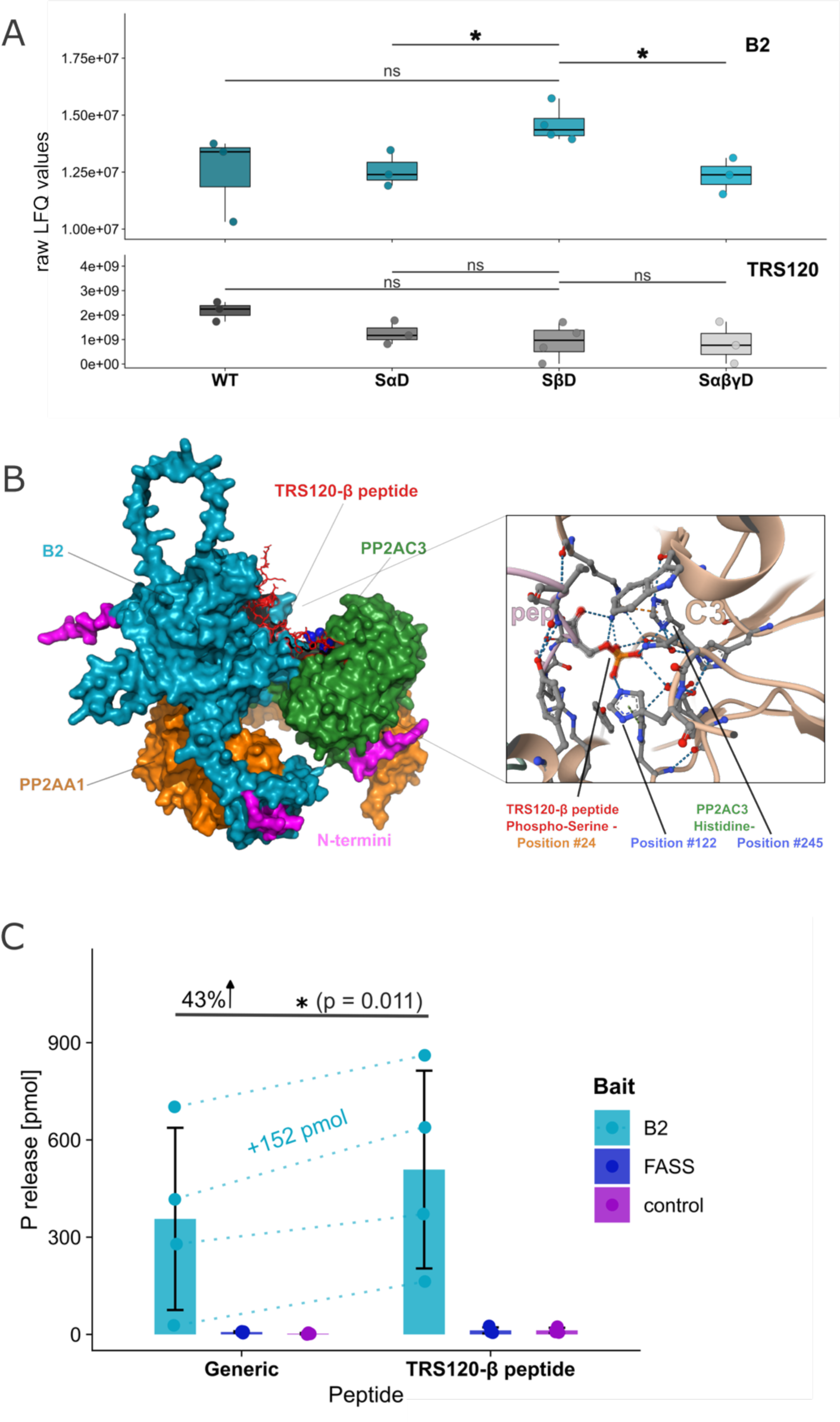
TRAPPII-PP2A interactions and establishment of an *in vitro* phosphatase assay. **A.** PP2A-B2 subunit enrichment in IP-MS of phosphomimetic TRS120 phosphovariants. Seedlings expressing AtTRS120-GFP with S-to-D substitutions in the phosphosite clusters were used (WT, SaD, SbD, and SabgD). The y-axis represents raw LFQ values for B2 (upper panel; note increased B2 intensity with TRS120-SβD) and TRS120 (lower panel; no significant difference in intensity) across TRS120 phosphovariants. Note that TRS120-SβD differed from SαD and SαβγD but not from WT. Three or four biological replicates were analysed for each phosphovariant. All values used are values from real IDs from the MS (no implementation of missing values). Protein abundance differences between genotypes were evaluated using pairwise two-sample t-tests, selecting Welch’s test when an F-test indicated unequal variances and Student’s t-test otherwise, with Benjamini–Hochberg FDR correction applied to adjust p-values across the three planned comparisons. n.s.: not significant. *: p < 0.05 **B.** AlphaFold 3 (Abramson et al., 2024) and AlphaBridge (Álvarez-Salmoral et al., 2024; inset, right) predictions of PP2A bound to the TRS120-β peptide substrate (EHFKLPVLDGSFFTKDPPPG[pSer]**P**S[pSer]SRNPSF; stick visualisation, red) with PP2AB2 in cyan, PP2AC3 in green, and PP2AA1 in orange. N-termini used for protein tagging are highlighted in magenta. Left: The TRS120-β peptide stretches over the surface of the regulatory B2 subunit with its phosphorylated residues (pS21, pS24) positioned near the catalytic centre of the C3 subunit (blue bubble visualisation). Inset, right: likely contact sites (dotted lines) between the phosphorylated Ser24 of the TRS120-β peptide (orange) and C3 histidines (residues 122 and 245; blue). See Fig. S4. **C.** *In vitro* phosphatase assay. Two different PP2A subunit combinations were transiently expressed, together with the PTPA chaperone (Chen *et al*., 2015), in *Nicotiana benthamiana* leaves for FLAG-IP. Control: FLAG-PUX5, a protein involved in ubiquitin regulation (Park et al., 2007), was used as a negative control for co-expression with PP2A A and C subunits (see Table S3). The y-axis shows the phosphate released (malachite green assay). Means ± SD. Jittered points represent the mean of three technical replicates for each of four independent biological experiments; dashed lines indicate within-experiment comparisons for the B2-FLAG IP. The x-axis shows the synthetic phosphopeptides used. Note that B2-FLAG preparations exhibited phosphatase activity towards the TRS120-β peptide. p-values were computed with a paired t-test (*: p < 0.05). Related to Fig. S5.

Structural modelling and interface analysis (Abramson et al., 2024; Salter et al., 2023; Álvarez-Salmoral et al., 2024) produced a model of an A1–B2–C3 PP2A complex with canonical PP2A architecture (Fig. S4). The highest-confidence interfaces linked A1 to both B2 and C3, consistent with the role of A1 as the structural scaffold for holoenzyme assembly (Fig. S4A). When a phosphorylated TRS120 peptide was included in the modelling approach, it was positioned at the interface between the B2 regulatory and C3 catalytic subunits (Fig. 5B; Fig. S4B). The predicted binding geometry was consistent with aspects of the catalytic arrangement proposed by previous quantum-based modelling of PP2A (Salter et al., 2023). The sequence context surrounding the TRS120-β site may be particularly relevant given the reported recognition of proline-containing phosphosites by B55-type PP2A complexes in fission yeast (Zeisner et al., 2025). PP2A B subunits mediate the trimeric complex’s subcellular localisation and functional regulation (Fahs et al., 2016; Hein et al., 2023). However, while B subunits have been documented to confer substrate specificity in a few cases in yeast (PP2A/Cdc55^reg^; Hoermann et al., 2020; Zeisner et al., 2025), and humans (PP2A-B55 and PP2A-B56; Hertz et al., 2016; Padi et al., 2024; Scheinost et al., 2026), the mapping of specific substrates to PP2A trimers harbouring specific B subunits remains poorly characterised in plants.

To test whether PP2A trimeric complexes containing the B2 regulatory subunit target the AtTRS120-β phosphosite cluster, we established a plant-based *in vitro* PP2A dephosphorylation assay. The assay was designed to assess the activity of plant PP2A holoenzymes on synthetic phosphorylated peptides. To reconstitute defined PP2A trimers, we heterologously expressed selected *Arabidopsis thaliana* PP2A subunits in leaves of *Nicotiana benthamiana* (Goodin et al., 2008). Given the considerable expansion of the PP2A family in Arabidopsis (Fig. S1A), there are theoretically 255 possible PP2A heterotrimers. However, which of these combinations exist *in vivo* remains to be determined. To address this for a subset of possible combinations, we assembled the catalytic subunit PP2AC3, the scaffolding subunit PP2AA1, a regulatory B subunit, together with the PP2A-activating chaperone PTPA (Chen et al., 2014). PTPA has been shown to promote the assembly of the A and C subunits and thereby increase PP2A activity (Chen et al., 2014; Chen et al., 2015). The four components were assembled on a single binary vector in which each gene was driven by a 35S promoter (Amack and Antunes, 2020; Fig. S5). Two of the 17 B subunits were chosen: B2, as it was differentially enriched in IP-MS with phosphomimetic AtTRS120-SβD compared with the other two phosphomimetics tested (Fig. 5A), and FASS as a representative of the B72 clade (Fig. S1A). Because the B subunit represents the gatekeeper of the catalytic cleft (Kruse et al., 2024; Fig. 5B), purifying PP2A via the regulatory B subunits may allow one to probe how trimer compositions with different classes of B subunits influence substrate selectivity.

For affinity purification, each B subunit was FLAG-tagged (Hopp et al., 1988) for mild competitive elution with FLAG peptide (Fig. S5A, S5B). We assessed complex assembly by co-immunopurification. FLAG-IP of B2 and FASS recovered all co-expressed subunits (Fig. S5C). These experiments demonstrate that, as predicted (Fig. S4), B2 associates with PP2AA1 and PP2AC3 following heterologous co-expression in *N. benthamiana*. As a substrate for the phosphatase assay, we used a synthetic 30-mer phosphopeptide encompassing two AtTRS120-β phosphoserines previously validated by *in vivo* phosphoproteomic and *in vitro* kinase assays (Mergner et al., 2020; Wiese et al., 2024). We refer to this peptide as the TRS120-β peptide (Fig. 5B, 5C). A generic 6-mer phosphopeptide was used as a nonspecific PP2A substrate control (Deana et al., 1990; Fig. 5C). B2-FLAG preparations exhibited phosphatase activity towards the TRS120-β peptide *in vitro* (Fig. 5C). In contrast, FASS FLAG-IPs showed no detectable phosphatase activity towards either the generic or TRS120-β peptides (Fig. 5C). This observation supports the phosphorylated TRS120 β-site peptide as an *in vitro* PP2A substrate and implicates B2 as a candidate regulatory subunit that may contribute to its recognition *in vivo*.

### Limitations of the study

The phosphatase assays used a doubly phosphorylated TRS120 peptide, which does not reproduce the structural context, site accessibility, or additional interaction surfaces of intact TRS120. FASS-FLAG preparations displayed no activity towards the generic control, precluding a definitive comparison of B-subunit-dependent substrate specificity. Because the complexes were reconstituted in *N. benthamiana*, endogenous PP2A components or other co-purifying proteins may have contributed to complex assembly and/or the measured activity. Thus, the data demonstrate phosphatase activity associated with B2-FLAG preparations *in vitro* but do not establish dephosphorylation of TRS120 *in vivo*.

## CONCLUSIONS

Differential proteomic analysis of TRS120 phosphomimetic variants *in vivo* (Fig. 5A), structural modelling (Fig. 5B; Fig. S4), and an *in vitro* phosphatase assay (Fig. 5C) converge on a model in which a B2-containing PP2A holoenzyme recognises and dephosphorylates a phosphorylated peptide encompassing the TRS120 β-site cluster. This interpretation is consistent with the established contribution of PP2A regulatory B subunits to substrate selection and with evidence that phosphoprotein phosphatase specificity depends on both the local sequence surrounding the phosphosite and higher-order substrate–holoenzyme interactions (Hein et al., 2023; Hertz et al., 2016; Hoermann et al., 2020; Kruse et al., 2024). In the structural model, the phosphorylated TRS120 β peptide spans the surface of B2 and is positioned at the B2-C3 interface, with its phosphoserines adjacent to the C3 catalytic centre. This arrangement raises the possibility that B2 contributes to recruiting and positioning the TRS120 β-site region for dephosphorylation by C3 (Fig. 5B; Fig. S4; Salter et al., 2023).

Taken together, our findings suggest that TRS120 phosphorylation regulates its engagement with a PP2A holoenzyme while, in turn, PP2A may contribute to regulating the TRAPPII phosphorylation status. Our findings further raise the possibility that PP2A counterbalances the SHAGGY-like kinase-mediated phosphorylation of TRS120 (Wiese et al., 2024). A direct convergence of SHAGGY-like kinases (AtSKs) and PP2A has already been demonstrated for mammalian PI3K-AKT-GSK-3β, where both target the same microtubule-associated protein TAU (Wang et al., 2015; Yao et al., 2011). Both SHAGGY-like kinases and PP2A holoenzymes function at multiple levels of plant signalling networks, including pathways initiated or regulated at the plasma membrane, and integrate diverse developmental, hormonal, biotic and abiotic cues (Máthé et al., 2026; Song et al., 2023). Their convergence on TRAPPII would provide a mechanistic link between signalling, membrane trafficking, the spatial control of plant development, and adaptation to environmental stress conditions.

## MATERIALS AND METHODS

### Lines and growth conditions

Arabidopsis mutant and transgenic lines are listed in Table S1. Plants were grown under controlled conditions as described in the Supplemental Materials and Methods. *Nicotiana benthamiana* was grown under long-day conditions for transient expression and PP2A purification.

### Yeast two-hybrid (Y2H) and molecular techniques

Pairwise Y2H assays were used to test interactions between TRS120 and CLUB/AtTRS130 truncations and selected PP2A ORFs (see Supplemental Materials and Methods). Interactions were assessed using the HIS3 reporter and considered positive only when reproducible in four independent assays. Phosphomimetic TRS120 phosphosite variants (TRS120-SαD, -SβD, and -SαβγD) were generated by site-directed mutagenesis as described (Wiese et al., 2024; Table S2).

### PP2A purification and phosphatase assay

PP2AA1, PP2A regulatory B-subunit, and PP2AC3 cDNAs, as well as a genomic PTPA construct, were transiently co-expressed in *N. benthamiana* (Table S3). B-subunit-containing complexes were affinity-purified via FLAG tags fused to the B subunits (B2, FASS) and competitively eluted with FLAG peptide. Purified complexes were tested against synthetic phosphopeptides, including a generic 6-mer (sequence: RRA[pThr]VA) and a 30-mer spanning the TRS120 β-site (sequence: EHFKLPVLDGSFFTKDPPPG[pSer]PS[pSer]SRNPSF), using a malachite green phosphate-release assay. Activity was compared with a control phosphopeptide and corrected using enzyme-only and peptide-specific controls. Detailed purification and assay conditions are provided in the Supplemental Materials and Methods.

### Light and electron microscopy

Scanning electron microscopy was used to assess differential growth phenotypes. Confocal microscopy was carried out on 3-day-old hypocotyls and 5-day-old root tips of plate-grown seedlings. Membrane association was quantified as the fluorescence ratio between membrane and matched cytoplasmic regions of interest. Cortical microtubules were immunolabelled with anti-α-tubulin and analysed after image segmentation and extraction of the cortical layer of epidermal cells. Microtubule organisation was quantified using FibrilTool-derived anisotropy measurements. Light-induced microtubule reorientation was additionally assessed in dark-grown seedlings by imaging the TUA5-mCherry marker (Gutierrez et al., 2009) in *trappii* mutants. For further details, see the Supplemental Materials.

### Modelling

AlphaFold 3 (Abramson et al., 2024) was used to model PP2A-TRS120 phosphopeptide complexes. AlphaBridge (Álvarez-Salmoral et al., 2024) was used to identify high-confidence inter-chain contacts and interfaces involving the phosphopeptide. See Supplemental Information.

### Mass spectrometry, LC-MS/MS data analysis and GO enrichment

TRAPPII-associated proteins were identified by GFP-based co-IP followed by label-free quantitative LC-MS/MS, as described by Wiese *et al*. (2024). Inflorescences were used for interactome analysis (Fig. 1), while 7-day-old seedlings were used to compare TRS120 phosphovariants (Fig. 5). Relative protein enrichment was determined in comparison to the empty vector GFP control for the pairwise comparisons (volcano plots in Fig. 1A and 1B), and proteins with ≥ 5-fold enrichment and P < 0.02 were considered significantly enriched. Gene Ontology enrichment was performed using clusterProfiler in R and the Arabidopsis TAIR annotation database. See Supplemental Materials.

### Statistical analysis

All data visualisations and statistical analyses were performed in RStudio and are described in the respective figure legends and the Supplemental Information.

## AUTHOR CONTRIBUTIONS

**Conceptualization:** FFA, DB, MP, ASM

**Methodology:** PFB, MeA, EF, AS, MiA, DB, MP, CL, ASM, CW

**Investigation:** FFA, MeA, EF, AS, MiA, MP, CL, ASM, CW, KB

**Visualization:** AS, MP, ASM

**Funding acquisition:** FFA, PFB, DB, CL

**Supervision:** FFA, PFB, EF, DB, MP, CL

**Writing - original draft:** FFA, ASM

**Writing - review & editing:** FFA, PFB, DB, MP, ASM, CL, CW, MiA

## ACKNOWLEDGMENTS

We thank Prof. Wilfried Schwab and members of his department for support. Thanks to Dr. Magalie Uyttewaal and Dr. Hakim Mireau for their useful suggestions. We are grateful to Yannik Schreckenberg, Aloïse Ducamp and Franziska Hackbarth for technical assistance and to Julia Hagn for curation of the proteomics data. We thank the WZW/TUM Centre for Advanced Light Microscopy (CALM), headed by Ramon Torres-Ruiz and Klaus Michel, for access to confocal microscopes. Thanks to Roman Meier at the TUMmesa facility, directed by Leonardo Teixeira and Balint Jakli, for supporting us with optimal growth conditions for our plants. This work has also benefited from the support of IJPB’s Plant Observatory platforms PO-Plants and PO-Cyto.

## DATA AVAILABILITY STATEMENT

The previously published CLUB–GFP IP–MS dataset is available through ProteomeXchange via PRIDE (Perez-Riverol et al., 2019) under accession PXD013016 (Kalde et al., 2019). The remaining mass spectrometry raw data, protein FASTA files and MaxQuant output will be deposited in PRIDE. All other data supporting the findings are included in the article and its Supplemental Information.

## SUPPLEMENTAL INFORMATION

**Figure S1.** The relative abundance of Arabidopsis PP2A subunits in the TRAPPII interactome

**Figure S2.** Cell lengths and widths of *trappii* and *pp2aa3* single and double mutants

**Figure S3.** Gene Ontology (GO) enrichment analysis of the TRS120 interactome

**Figure S4.** Predicted PP2A-TRS120 interactions at the phosphorylated TRS120-β phosphosite cluster

**Figure S5.** Purification of Arabidopsis PP2A complexes from *Nicotiana benthamiana*.

## Supplemental Materials and Methods

**Table S1.** Mutant lines used in this study

**Table S2.** Primer sequences used for site-directed mutagenesis for TRS120 phosphovariants

**Table S3.** Primer sequences used for domestication of PP2A subunits and PTPA for phosphatase activity assay

**Table S4.** piCSi output values from AlphaBridge analysis

