## Supplemental Information for "The PP2A phosphatase associates with the Arabidopsis TRAPPII tethering complex and dephosphorylates a TRAPPII-derived phosphopeptide *in vitro*"

**Figure S1.** The relative abundance of Arabidopsis PP2A subunits in the TRAPPII interactome

**Figure S2.** Cell lengths and widths of *trappii* and *pp2aa3* single and double mutants

**Figure S3.** Gene Ontology (GO) enrichment analysis of the TRS120 interactome

**Figure S4.** Predicted PP2A-TRS120 interactions at the phosphorylated TRS120-β phosphosite cluster

**Figure S5.** Purification of Arabidopsis PP2A complexes from *Nicotiana benthamiana*.

**Supplemental Materials and Methods**

**Table S1.** Mutant lines used in this study

**Table S2.** Primer sequences used for site-directed mutagenesis for TRS120 phosphovariants

**Table S3.** Primer sequences used for domestication of PP2A subunits and PTPA for phosphatase activity assay

**Table S4.** piCSi output values from AlphaBridge analysis

A

| PP2A subunits in *Arabidopsis thaliana* | | | | | | |
|---|---|---|---|---|---|---|
| **PP2A Family** | **Subclade** | **subunit No.** | **AGI Locus Identifier** | **Greek letter** | **Alternative Nomenclature** | **Used in this study** |
| **A scaffolding** | **PR65*** | **A1 / RCN1** | AT1G25490 | - | ROOTS CURL IN NAPHTHYLPHTHALAMIC ACID1, PR65A1 | as **PP2AA1** |
| | **PR65*** | A2 | AT3G25800 | - | PR65A2 | |
| | | **A3** | AT1G13320 | - | PR65A3 | as **PP2AA3** |
| **B regulatory** | **B55/PPP2R2** | B1 | AT1G51690 | Bα | B55α | |
| | | **B2** | AT1G17720 | Bβ | B55β | as **B2** |
| | **B56/PPP2R5** | B3 | AT5G03470 | B'α | B56α | |
| | | B4 | AT3G09880 | B'β | B56β | |
| | | B5 | AT4G15415 | B'γ | B56γ | |
| | | B6 | AT3G26030 | B'δ | B56δ | |
| | | B7 | AT3G54930 | B'ε | B56ε | |
| | | B8 | AT3G21650 | B'ζ | B56ζ | |
| | | B9 | AT3G26020 | B'η | B56η | |
| | | B10 | AT1G13460 | B'θ | B56θ | |
| | | B11 | AT5G25510 | B'κ | B56κ | |
| | **B72/PPP2R3** | **FASS** | AT5G18580 | FASS/TON2 | - | as **FASS** |
| | | B13 | AT5G44090 | B"α | - | |
| | | B14 | AT1G03960 | B"β | - | |
| | | B15 | AT1G54450 | B"γ | - | |
| | | B16 | AT5G28850 | B"ε | - | |
| | | B17 | AT5G28900 | B"δ | - | |
| **C catalytic** | **class I** | C1 | AT1G59830 | - | - | |
| | | C2 | AT1G10430 | - | - | |
| | | C5 | AT1G69960 | - | - | |
| | **class II** | **C3** | AT2G42500 | - | - | as **PP2AC3** |
| | | C4 | AT3G58500 | - | - | |

*no formal subfamily definition, but A1 is functionally very distinct from A2/A3 (Blakeslee et al. 2008)

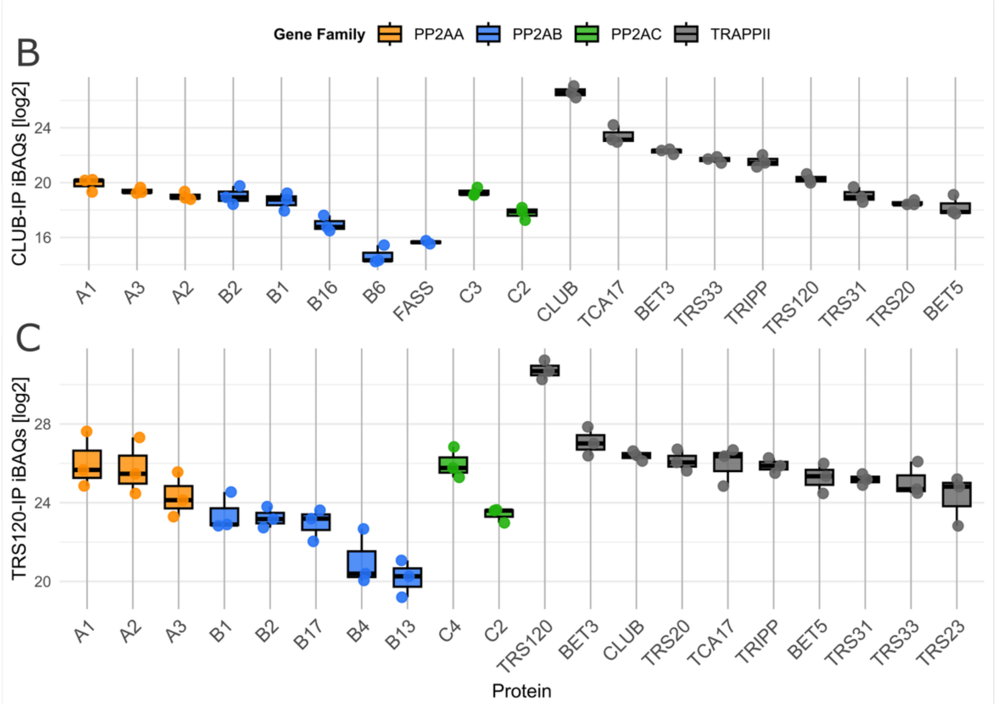

**Figure S1. The relative abundance of Arabidopsis PP2A subunits in the TRAPPII interactome.**
**A.** Nomenclature(s) of PP2A subunits in *Arabidopsis thaliana,* adapted from Booker & DeLong (2017). Isoforms used in this study are highlighted in bold.
**B-C.** PP2A subunit abundance in the CLUB-GFP (B) and TRS120-GFP (C) TRAPPII interactomes**.** Means ± SD of $log_2$ intensity-based absolute quantification (iBAQ values) of PP2A and TRAPPII subunits. The observed values from biological replicates (no imputation of missing values) are depicted with jittered points. iBAQ values were used to provide the closest approximation for relative molar abundance (Schwanhäusser et al., 2011). The CLUB and TRS120 baits were the most abundant proteins in the IP samples. The Arabidopsis PP2A subunits used to query the datasets are listed in panel A. Note that, among PP2A components, A subunits were most abundant. Within the B subunit family, B1 and B2 (B55 subfamily) were the most abundant. For catalytic subunits, both PP2AC subfamily II members (C3 in CLUB and C4 in TRS120) were detected. Related to Fig. 1A, B.

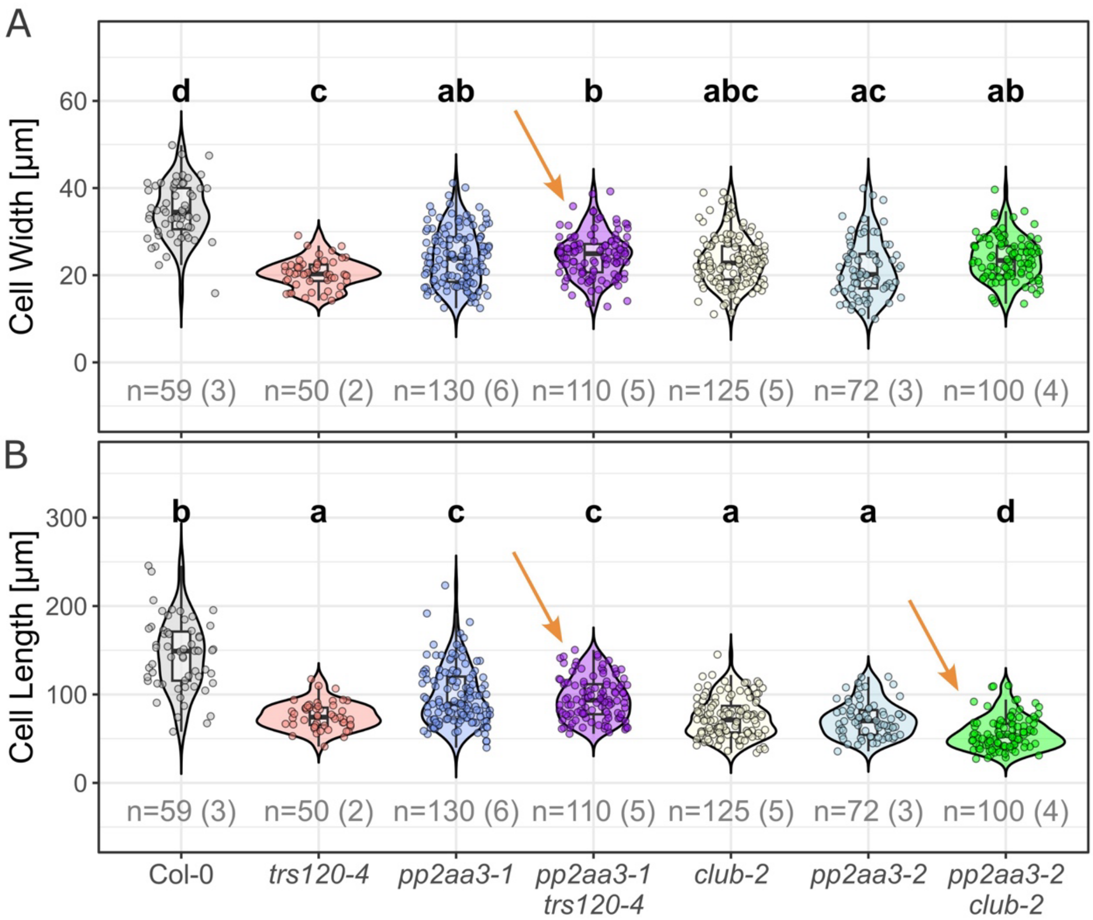

**Figure S2. Cell lengths and widths of *trappii* and *pp2aa3* single and double mutants.**
**A**. Hypocotyl cell widths in the epidermis; note that *trs120-4 pp2aa3*-1 double mutants had significantly wider cells than *trs120-4* (orange arrow).
**B.** Hypocotyl cell lengths in the epidermis; note that *trs120-4 pp2aa3*-1 double mutants had significantly longer cells than *trs120-4*; in contrast, *club-2 pp2aa3*-2 double mutants had significantly shorter cells (orange arrows).
P values were computed using Kruskal–Wallis followed by Bonferroni-corrected Dunn tests.
P values are represented by compact letter displays. Sample sizes (n) in grey = number of cells (number of seedlings). Related to Fig. 2.

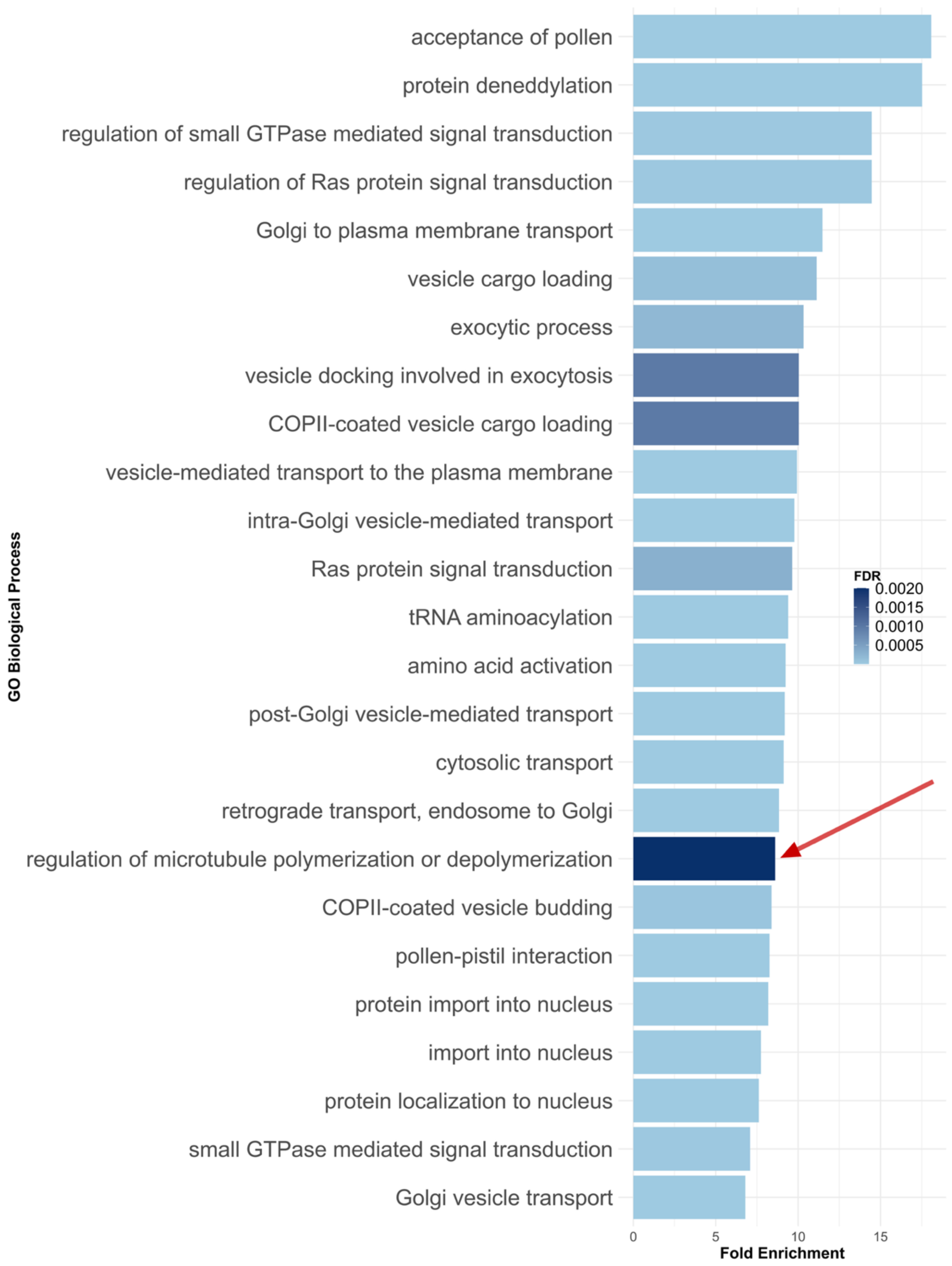

**Figure S3. Gene Ontology (GO) enrichment analysis of the TRS120 interactome.**
Proteins enriched in the TRS120 co-immunoprecipitation LFQ dataset were subjected to GO term enrichment analysis using the Biological Process (BP) ontology. Shown are significantly enriched GO terms (FDR ≤ 0.003) with strong over-representation (fold enrichment ≥ 6.5) after removal of broad housekeeping categories (biosynthetic, metabolic, translational, catabolic, glycolytic, pentose, gluconeogenesis, and oxidation-related terms). Bar length represents the fold enrichment of each GO term relative to the Arabidopsis genome background, while bar colour indicates statistical significance (FDR-adjusted p-value). GO terms are ordered by fold enrichment. This analysis highlights biological processes enriched among proteins associated with the TRAPPII-specific TRS120 subunit. Note that “regulation of microtubule polymerization or depolymerization” is among the most enriched terms (red arrow). Related to Fig. 3.

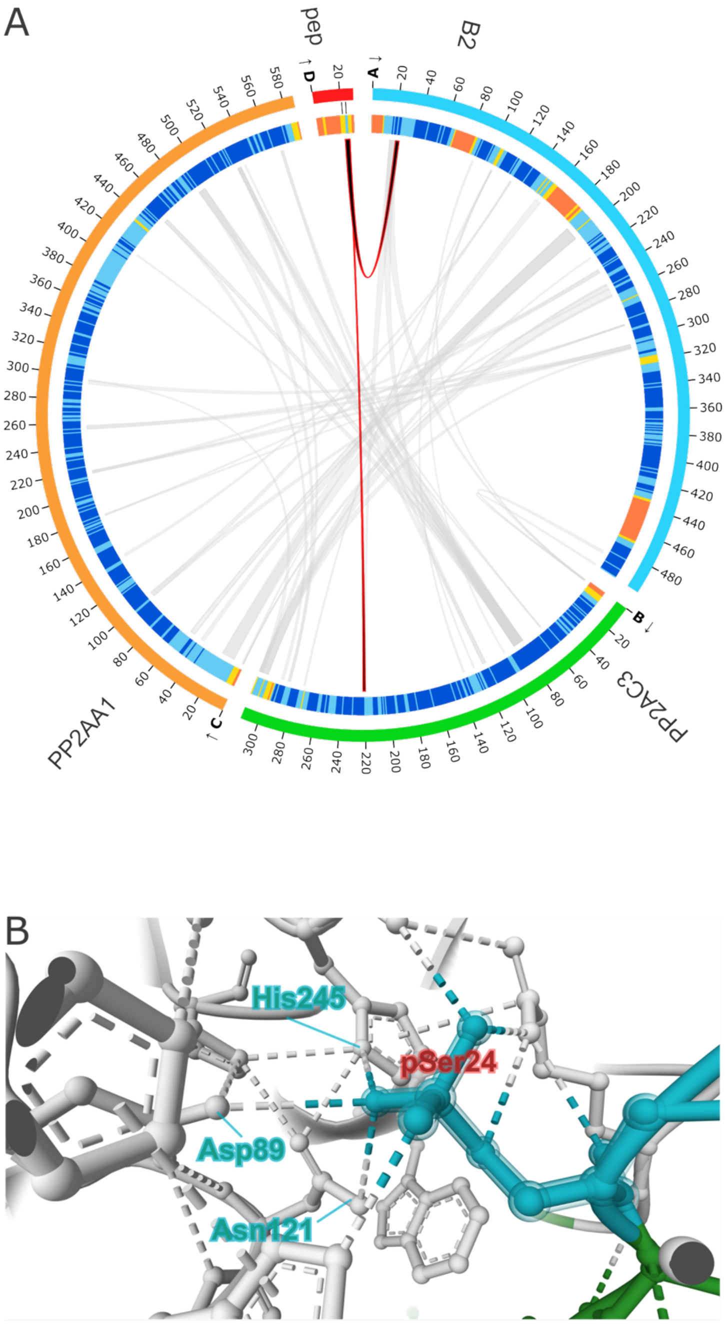

**Figure S4. Predicted PP2A-TRS120 interactions at the phosphorylated TRS120-β cluster.**
**A.** Protein–protein interfaces predicted by AlphaFold 3 (Abramson et al., 2024) and interpreted with AlphaBridge (Álvarez-Salmoral et al., 2024), using the default thresholds. Outer-ring colours indicate PP2AA1 (orange), B2 (cyan), PP2AC3 (green) and the TRS120-β phosphopeptide (red). Inner-ring colours indicate AlphaFold 3 pLDDT confidence scores. Two red links connect the phosphorylated residues of the TRS120-β peptide with catalytic PP2AC3 and regulatory B2. The strongest predicted interfaces connected B2 with the PP2A scaffold A1 (piCSi = 0.95; see Methods and Table S4) and C3 with A1 (piCSi = 0.96), whereas the B2–C3 contacts were weaker (piCSi = 0.84–0.86), supporting A1 as the principal structural platform for holoenzyme assembly.
**B.** Structural model and predicted contacts (related to Fig. S4A). Consistent with quantum-mechanical modelling of PP2A–substrate interactions (Salter et al., 2023), the model places one oxygen atom of the phosphate group of TRS120-β pSer24 in predicted contact with PP2AC3 residues Asp89, Asn121 and His245. Related to Fig. 5.

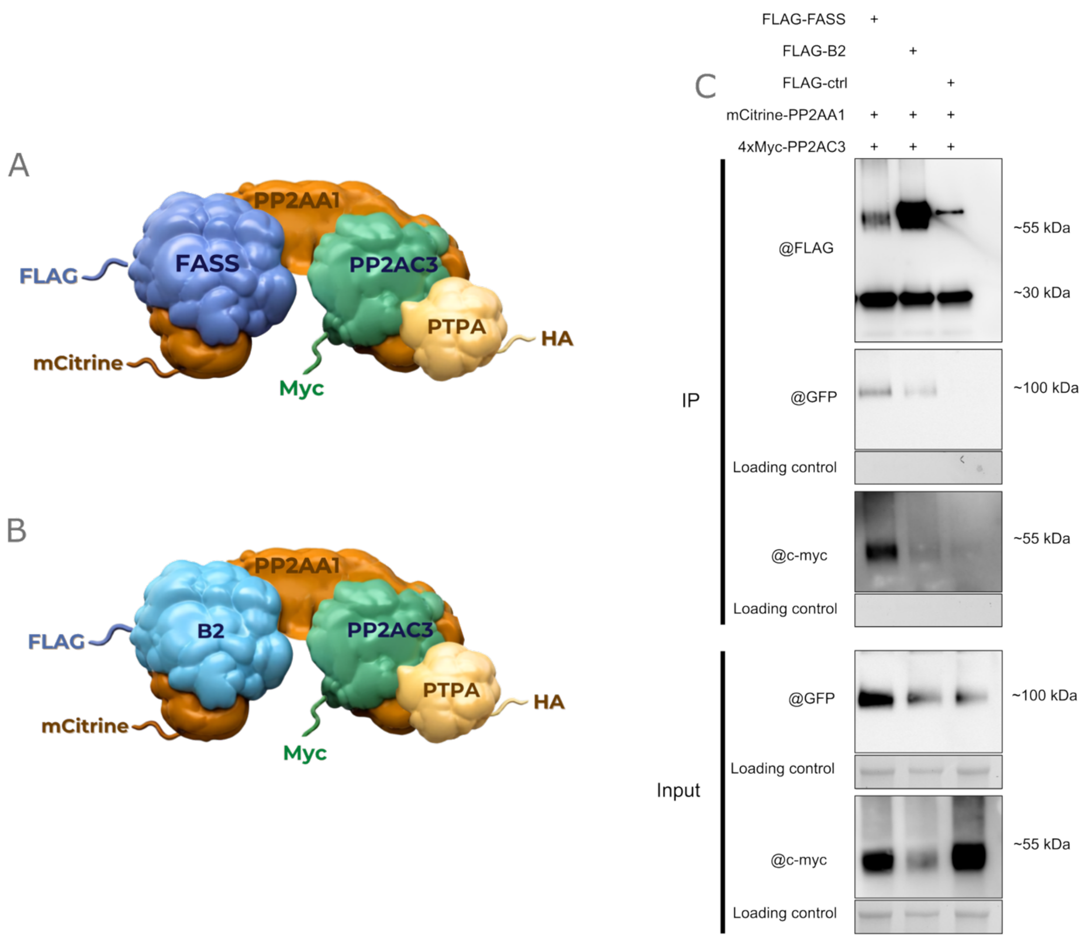

**Figure S5. Purification of Arabidopsis PP2A complexes from *Nicotiana benthamiana*.**

**A-B.** Schematics of the Arabidopsis PP2A complexes encoded by the multigene expression constructs. The mCitrine-PP2AA1 scaffolding subunit (orange), 4xMyc-PP2AC3 catalytic subunit (green) and chaperone 3xHA-PTPA (yellow) were identical in all constructs. In each expression construct, a different PP2A regulatory B subunit was FLAG-tagged for FLAG-IP.

**A. FLAG-FASS** (dark blue).

**B. FLAG-B2 (**light blue).

**C.** The FLAG-tagged bait proteins were visualised using anti-FLAG antibody. The other components were visualised with anti-GFP (for mCitrine-tagged PP2AA1) and anti-myc (for PP2AC3) antibodies. Total protein visualised using Bio-Rad Stain-Free technology served as a loading control. FLAG immunoprecipitation recovered PP2AA1 and PP2AC3 with FASS and B2, although partner recovery differed between preparations. In the control (ctrl; FLAG-PUX5) preparation, PP2AC3 was detected only weakly and PP2AA1 was not detected, indicating a specific recovery of the B subunits B2 and FASS. These experiments demonstrate that B2 associates with PP2AA1 and/or PP2AC3. Bands were detected at the expected sizes of: FLAG-FASS: 56 kDa; FLAG-B2: 57.3 kDa; mCitrine-PP2AA1: 92.6 kDa; 4xMyc-PP2AC3: 40.6 kDa; apparent molecular masses were estimated using protein ladders (relevant ladder sizes indicated on the right). Related to Fig. 5.

### SUPPLEMENTAL MATERIALS AND METHODS

#### Lines and growth conditions

All the mutant lines used in this study are listed in Table S1. Seedling-lethal mutants were propagated as hetero- or hemi-zygotes. Insertion lines were selected via the TAIR and NASC websites (Swarbreck et al., 2008). Plants for the Co-IP-MS were grown in the greenhouse under controlled temperature conditions with supplemental light, or in a growth chamber at the TUMmesa ecotron (16/8 h photoperiod at 180 µmol $m^{-2}s^{-1}$). Seeds were surface sterilised, stratified at 4 °C for 2 days, and plated on ½ MS medium supplemented with B5 vitamins (G1019; Sigma-Aldrich). For confocal microscopy, the medium was additionally supplemented with 1% sucrose. Plates were incubated at 22 °C under constant light (80 µmol $m^{-2}s^{-1}$). The root tips of 5-day-old plate-grown seedlings were used for confocal microscopy. 7-day-old plate-grown seedlings or inflorescences were used for co-immunoprecipitation.

For transient expression, *N. benthamiana* plants were grown under long-day conditions (16 h light/8 h dark; day: 24 °C and 40% relative humidity; night: 18 °C and 55% relative humidity), with 1-h transitions between regimes.

#### Molecular techniques and site-directed mutagenesis (SDM)

Standard molecular techniques were used for subcloning (Sambrook et al., 1989). RIKEN cDNA clones (Seki et al., 2002) served as templates for constructs used for expression in *E. coli*, *S. cerevisiae* or *N. benthamiana* (Beritza et al., 2024; Cordier et al., 1999). Phosphomimetic TRS120 phosphosite variants (SαD, SβD, SαβγD; see Table S2; Wiese et al., 2024) were generated using a DpnI-mediated site-directed mutagenesis protocol and the Gateway cloning system. Briefly, site-directed mutations were introduced into the template construct via polymerase chain reaction using mutagenic primers with the desired mutations (see Table S2) and the KOD Hot Start DNA Polymerase (Novagen) for strand extension. Subsequently, the methylated non-mutated DNA template was digested with the DpnI endonuclease (Thermo Fisher Scientific). Mutated vectors were transformed in E. coli DH5α for nick repair and amplification of the plasmids.

Following plasmid purification, constructs were sequence-verified. Variants containing mutations at multiple phosphosite clusters were generated by sequential rounds of mutagenesis using previously mutated plasmids as templates. For *in planta* experiments in Arabidopsis, such as IP-MS, the genomic construct ProTRS120:TRS120-GFP in the pCAMBIA2300 plasmid (Rybak et al., 2014) was used as an SDM template. Constructs were transformed into the *Agrobacterium tumefaciens* strain GV3101(pMP90) and introduced into the Arabidopsis hemizygous null *trs120-4* background using the floral dip method (Clough and

Bent, 1998). Transgenic lines were selected on ½ MS medium supplemented with 50 μg/ml kanamycin. For live-cell confocal imaging of the PP2AA subunits, we used ProPP2AA1:YFP-PP2AA1, ProPP2AA3:YFP-PP2AA3 (Blakeslee et al., 2008) in segregating *trs120-4* lines.

**Yeast two-hybrid (Y2H)**

Y2H pairwise tests were performed as described in (Altmann et al., 2020). Briefly, cDNA fragments encoding CLUB/AtTRS130 truncations C1–C3 and TRS120 truncations T1 and T3 (Kalde et al., 2019; Wiese et al., 2024) were transferred by Gateway cloning into the GAL4 DNA-binding-domain vector pDEST-pPC97 and transformed into yeast strain Y8930. These strains were mated with Y8800 strains carrying GAL4 activation-domain fusions of PP2AA1 (At1g25490), PP2AA2 (At3g25800), PP2AA3 (At1g13320), TON2/FASS (At5g18580), PP2AC3 (At2g42500), PP2AC4 (At3g58500). Interactions were assayed using the HIS3 reporter on medium containing 1 or 5 mM 3-AT; results for 1 mM are shown. All candidate interactions were verified by pairwise one-on-one mating in four independent experiments. Only pairs that scored positively in all four assays were considered reproducible interaction partners. The use of low-copy plasmids, weak promoters, the counter-selectable marker cyh2S on the AD-Y plasmid, as well as semi-quantitative scoring of quadruplicate tests has been shown to reliably eliminate experimental artefacts and hence false positives. With the exception of the CLUB-C1 truncation, all TRAPPII truncations yielded at least one positive interaction in pairwise tests (see also Kalde et al., 2019; Garcia et al., 2020), and this was used as an internal positive control for the interpretation of negative interaction data.

**Transient expression of Arabidopsis PP2A subunits in *Nicotiana benthamiana***

Two to three fully expanded leaves from 3- to 4-week-old wild-type *N. benthamiana* plants at the eight-leaf stage were co-infiltrated with *Agrobacterium tumefaciens* strains containing the expression plasmids with the PP2A subunits and the p19 helper plasmid (Jay et al., 2023). The plasmids were generated using the GoldenBraid (Sarrion-Perdigones et al., 2013) system by cloning full-length cDNAs encoding PP2AA1, PP2AC3, B2, FASS, and PUX5, while PTPA was amplified from genomic DNA (Col-0). Epitope tags (3xFLAG, 4xMYC, 3xHA, and mCitrine) were incorporated as modular fusion elements at the N terminus for heterologous expression. All expression units were assembled on the same vector. Infiltration was performed with Agrobacterium cultures carrying the p19 helper plasmid or expression vectors, each adjusted to OD600 = 0.5 and mixed in equal volumes, in transformation medium (Duchefa S-Medium supplemented with 4% sucrose, pH 5.7). Infiltrated areas were cut out 3 days post-infiltration, snap-frozen in liquid nitrogen and stored at −80 °C until further processing.

**Enzyme purification**

1.5 g of frozen *Nicotiana benthamiana* leaf tissue was ground to a powder in liquid nitrogen and resuspended in 1.5 mL of IP buffer (adapted from Skottke et al., 2011; final composition: 50 mM Tris-HCl, pH 7.5; 100 mM NaCl; 0.3 M sucrose; 0.1% (v/v) Triton X-100; 5% (v/v) glycerol; Protease inhibitor cocktail (Sigma fast EDTA free®); 1 mM DTT; 1mM PMSF (phenylmethylsulfonyl fluoride)). Samples were rotated for 90 min at 4 °C. Lysates were centrifuged for 60 min at 20,238 × g at 4 °C. The supernatants (approximately 3 mL) were incubated with 125 µL anti-FLAG-Agarose bead slurry (Chromotek®) for 90 min at 4 °C. Beads were washed 3 times in 1 mL IP buffer and then eluted with 240 µL peptide elution buffer (50 mM Tris-HCl pH 7.5, 100 mM NaCl, 0.3 M sucrose, 5% glycerol, Protease inhibitor cocktail (Sigma fast EDTA free®), 1 mM DTT, 300 µg/mL 3×FLAG peptide (Chromotek®)) for 30 min at room temperature. Eluates were briefly stored on ice until use in the enzyme assays.

**PP2A assay**

Reactions were mixed directly, quickly, and thoroughly in clear 96-well plates with half-area (Corning®). Each 50 µL reaction contained 10 µL of 5× PP2A reaction buffer (250 mM imidazole, pH 7.2, 1 mM DTT, 0.5 mg $mL^{-1}$ BSA, 1 mM $MnCl_2$ and 1 mM $CaCl_2$; final concentrations: 50 mM imidazole, 0.2 mM DTT, 0.1 mg $mL^{-1}$ BSA, 0.2 mM $MnCl_2$ and 0.2 mM $CaCl_2$; modified from the Promega Serine/Threonine Phosphatase Assay), 20 µL of eluate containing purified enzyme, 10 µL of 1 mM synthetic phosphopeptide (200 µM final; GenScript), and 10 µL of ultrapure water. The plate was incubated at 37 °C for 60 min in the dark. The reactions were terminated by adding 50 µL of malachite green reagent from the Phosphate Assay Kit (MAK307; Sigma-Aldrich). The absorbance at 600 nm was measured after a 30 min incubation with the reagent at room temperature. The enzyme assay for each biological replicate was performed in three technical replicates, and the means of the technical replicates were used as the value for the biological replicate. Phosphatase activity values were derived from in-plate phosphate standards. For each bait and biological replicate, activity was corrected by subtracting the enzyme-only control. Values for the TRS120-β peptide were additionally corrected for peptide-specific background measured in no-enzyme controls. Generic (RRA[pThr]VA) and TRS120-β peptide (EHFKLPVLDGSFFTKDPPPG[pSer]PS[pSer]SRNPSF) activities were compared within biological replicates using a paired, two-sided t test. Mean paired increase was calculated as the mean of the within-replicate differences and is reported in pmol phosphate.

**Light and electron microscopy**

For scanning electron microscopy (SEM), a Zeiss (LEO) VP 438 microscope was operated at 15 kV. 10-day-old seedlings from the differential growth decision assay were placed onto stubs and examined immediately in low vacuum.

Confocal microscopy imaging was performed at a controlled room temperature (22°C) using a Leica (https://www.leica-microsystems.com) TCS SP8 X HyVolution CSLM with a 40× 1.1 NA or 63× 1.2 NA water-immersion objective. Imaging data were acquired using LAS X software (Leica). The live-imaging data were quantified manually using FIJI (Schindelin et al., 2012). For the SEMs the line tool from ImageJ was drawn from approximately the middle of each side to the opposing side. For quantification of membrane association, images were acquired at the focal plane in which the nucleus was in focus and the cell boundaries were clearly resolved. For each cell, equal-sized rectangular ROIs were placed in paired positions within the same cell: one ROI was positioned over the plasma membrane, and the corresponding ROI was positioned in the adjacent cytoplasm at the same distance from the nucleus. The same ROI dimensions were used for all measurements. Mean fluorescence intensity was measured for each ROI, and membrane association was expressed as the ratio of membrane to cytoplasmic fluorescence intensity. Background fluorescence was subtracted from all measurements before calculation of the ratio. Images used for quantification were acquired using an identical microscope, objective, laser power, detector gain, exposure, and other acquisition settings for all experimental conditions. P-values were computed with a Wilcoxon rank-sum test in R.

**Immunostaining and semiquantitative analysis of cortical microtubules**

4-day-old seedlings were fixed in 4% paraformaldehyde and 1.5 % Triton X-100 in 0.5× MTSB buffer (25 mM PIPES, 2.5 mM $MgSO_4$, 2.5 mM EGTA, pH 6.9) for 1 h under vacuum, then rinsed in 1× PBS for 10 min. Cell walls were then digested using the following buffer for one hour: 2 mM MES pH 5, 0.2 % driselase and 0.15 % macerozyme. Tissues were incubated overnight at room temperature with the B-5-1-2 monoclonal anti-α-tubulin antibody (T9026; Sigma-Aldrich; 1:2,000). The next day, tissues were washed for 20 min in PBS containing 50 mM glycine and incubated with the Alexa Fluor 555 goat anti-mouse secondary antibody (A21422; Life Technologies, Carlsbad, CA, USA; 1:2,000) over-night and washed again in PBS containing 50 mM glycine. Samples were then incubated for 30 min in 0.01 % Calcofluor (Fluorescent Brightener 28) in PBS, then rinsed in PBS before mounting in the Vectashield (Vector Laboratories, Inc., Newark, USA) mounting medium.

Samples were viewed using a Stellaris8 STED-FLIM confocal microscope (Leica Microsystems). Roots were excited sequentially at 405 nm (Calcofluor) and 553 nm (Alexa fluor 555) with an emission band of 430-480 nm (Calcofluor) and 560-620 nm (Alexa fluor 555). All stacks were imaged using the same zoom (x 2.25) with voxel dimensions of 80 × 80 × 300 nm.

The calcofluor channel was denoised using a 3D-Gaussian Blur filter (sigma = 1). Images were resized and submitted to the PlantSeg pipeline using the generic_confocal_3D_unet model (Wolny et al., 2020). The resulting prediction image was resized to the original size and segmented with the BIP software (h-watershed algorithm, h-value = 20). Epidermal cells were extracted using the MorphoLibJ plugin (Legland et al., 2016). Dividing cells were excluded from the analysis. The SurfCut ImageJ macro (Erguvan et al., 2019) was used to extract a layer of 2.5 µm depth from all labels, corresponding to the cortical zone of the external faces of epidermal cells. This stack resulting from the Surfcut process was then binarized and multiplied by the microtubules stack.

A maximum projection was generated for each SurfCut label stack. Labels were then eroded with a radius of 7 using the MorphoLibJ Label Morphological Filters module and converted into Fiji Regions Of Interest (ROI) via the MorphoLibJ Label Map to ROIs module. This erosion step excluded microtubule signals at cell contours from subsequent analysis. The FibrilTool batch version of the FibrilTool Fiji plugin was used (Boudaoud et al., 2014) to compute automatically the anisotropy level of microtubule arrays in the set of ROIs.

**Microtubule reorientation assay**

For the microtubule reorientation assay (Kirik et al., 2012), 3-day-old dark-grown wild-type and *trappii* (*club-2, trs120-4*) seedlings were exposed to light using the incandescent light source that was aligned for Köhler illumination on the FV1000 (Olympus) confocal microscope at full power. Undamaged hypocotyl cells were selected. Microtubules were imaged using TUA5-mCherry (Gutierrez et al., 2009), and their orientation angles were scored into 10° bins by tracing with the line tool and cell counter plugin in ImageJ.

**Modelling**

AlphaFold 3 (Abramson et al., 2024; https://alphafoldserver.com/) was used to generate multimeric protein models from TAIR sequences and sequences of our synthetic phosphopeptide, producing five structural samples per complex together with pLDDT, PAE, PDE, and interface confidence metrics; the highest-confidence model was selected based on global pLDDT and inter-chain PAE distributions (model 0). Interfaces were defined using AlphaBridge's default custom contact-probability cutoff (0.7), and the piCSi confidence score was then computed by AlphaBridge per interface from high-confidence contact probabilities (CP) and normalized PMC-derived structural confidence (CPMC) (Álvarez-Salmoral et al., 2024; https://alpha-bridge.eu/ - used on 20/08/2026). Structural inspection and visualisation were performed in PyMOL (Schrödinger, LLC).

### Mass spectrometry-based Proteomics

For TRAPPII bait proteins, co-immunoprecipitation experiments were performed using 3 g of tissue and GFP-Trap beads (Chromotek©), following the procedure described previously (Kalde et al., 2019, Rybak et al., 2014; Wiese et al. 2024).

In-gel digestion with trypsin was conducted according to established procedures (Shevchenko et al., 2006). Briefly, samples were separated on a NuPAGE™ 4 - 12% Bis-Tris protein gel (Thermo Fisher Scientific) for approximately 1 cm. The resulting, still largely unresolved single protein band from each sample was excised and subjected to reduction with 10 mM dithiothreitol in 5 mM TEAB, followed by alkylation with 55 mM chloroacetamide in 5 mM TEAB. Samples were then digested overnight with mass-spectrometry-grade Trypsin Gold (Promega) for 1h at 30°C in a ratio of 1:100 trypsin: protein. After 1h incubation, the same amount of trypsin was added a second time. The reaction was stopped with 1% formic acid after 16 h. The resulting peptides were dried completely and subsequently resuspended in 24 µL of 2% acetonitrile and 0.1% formic acid in HPLC-grade water. For each mass spectrometric measurement, 10 µL of the peptide solution was injected.

### LC-MS/MS data acquisition

LC-MS/MS measurements were performed using a Dionex UltiMate 3000 RSLCnano system coupled to an Orbitrap Fusion LUMOS mass spectrometer (Thermo Fisher Scientific, Bremen). Peptide samples were initially loaded onto a trap column consisting of ReproSil-pur C18-AQ, 5 µm particle size (Dr. Maisch; 20 mm × 75 µm; self-packed), at a flow rate of 5 µL/min using 0.1% formic acid in HPLC-grade water. Following a 10-min loading period, peptides were transferred to a self-packed analytical column containing ReproSil Gold C18-AQ, 3 µm particle size (Dr. Maisch; 450 mm × 75 µm) over 1 min. Peptides were subsequently separated at a flow rate of 300 nL/min using a 50-min gradient from 4% to 32% solvent B. Solvent B consisted of 0.1% formic acid and 5% DMSO in acetonitrile, whereas solvent A contained 0.1% formic acid and 5% DMSO in HPLC-grade water.

The Orbitrap Fusion LUMOS was operated in positive ionization mode using data-dependent acquisition (DDA). MS1 spectra were acquired over an m/z range of 360–1300 at a resolution of 60,000, with an automatic gain control (AGC) target of 4 × 10^5 and a maximum injection time (maxIT) of 50 ms. The acquisition cycle time was set to 2 s. Precursors carrying charge states of 2–6 were selected for fragmentation, and dynamic exclusion was applied for 30 s. Peptide fragmentation was performed by higher-energy collisional dissociation (HCD) using a normalized collision energy (NCE) of 30%. The precursor isolation window was set to 1.3 m/z. MS2 spectra were acquired at a resolution of 15,000 with an AGC target of 7.5 × 10^4 and a maximum injection time of 22 ms.

All proteomic samples were measured in randomized order. A blank injection was performed between consecutive samples to permit column re-equilibration and reduce sample carryover. Instrument and overall system performance were monitored throughout the measurements by regular injections of a Pierce HeLa digest quality-control sample.

**LC-MS/MS data analysis**

Peptide identification and quantification were performed using MaxQuant software (version 2.4.8.0). MS2 spectra were searched against the Arabidopsis thaliana protein database obtained from The Arabidopsis Information Resource (TAIR) with common contaminants included using the corresponding built-in MaxQuant option. Trypsin/P was specified as the proteolytic enzyme with up to two missed cleavage sites. Carbamidomethylation of cysteine was defined as a fixed modification, while oxidation of methionine and acetylation of protein N-termini were specified as variable modifications.

A false discovery rate (FDR) of 1% was applied at both the peptide-spectrum match (PSM) and protein levels using a target-decoy approach based on reversed protein sequences. Protein quantification was performed using both label-free quantification (LFQ) and intensity-based absolute quantification (iBAQ), requiring a minimum of two peptides per protein. The minimum peptide length was set to seven amino acids. The “match between runs” function was enabled using a matching time window of 0.4 min and an alignment window of 20 min.

The resulting protein table was subsequently analysed using the Perseus data analysis program (v2.1.2.0). Data processing included filtering for proteins detected in at least two biological replicates in at least one condition, followed by imputation of missing values from a normal distribution (width = 0.3; downshift = 1.8). Differential protein abundance between Co-IP samples from EGAD-GFP and either TRS120-GFP WT or CLUB-GFP WT was assessed using Welch’s two-tailed t-test.

Proteins showing a fold change of at least 5 and a P-value below 0.02 were considered significantly enriched.

**GO enrichment**

Gene Ontology enrichment analysis was performed using the clusterProfiler 4.0 package in R (Wu et al., 2021) to identify over-represented Biological Process terms associated with the TRAPPII interactome. Arabidopsis TAIR locus identifiers (AGIs) extracted from the proteomics dataset of EGAD-GFP vs TRS120-GFP WT with at least twofold enrichment in TRS120-GFP relative to EGAD-GFP (974 of 2,311 identified proteins were retained) were used as input for enrichment against the org.At.tair.db annotation database (version 3.18.0). Over-representation analysis was performed using enrichGO, with TAIR IDs as the key type, the *Biological Process* ontology, and Benjamini–Hochberg correction for multiple testing. Terms were considered significantly enriched at q value < 0.05. Fold enrichment was calculated from the GeneRatio and BgRatio values, and terms were subsequently filtered for fold enrichment ≥ 6.5 and FDR ≤ 0.003. Terms associated with broad biosynthetic, metabolic, translational, catabolic, gluconeogenic, pentose, and glycolytic processes were excluded from downstream interpretation.

**Statistical analysis and image processing**

Raw numerical data were recorded in Excel. Subsequent data analysis, statistical testing, and visualization were performed in RStudio (version 2026.7.0.139) and R (R Core Team, 2021; version 4.6.1). All datasets were first inspected for distributional properties and homoscedasticity using visual diagnostics (violin/boxplots and jittered raw data). For comparisons involving multiple groups, approximately symmetric data were analysed using a one-way ANOVA followed by Tukey’s HSD for all pairwise comparisons. Measurements that showed clear deviations from normality and unequal variances were analysed using a non-parametric Kruskal–Wallis test followed by Dunn’s post-hoc tests with Bonferroni correction to control the family-wise error rate across multiple comparisons. The respective statistical tests for each dataset are indicated in the figure legends.

**Table S1. Mutant lines used in this study**

| Allele | AGI locus | Polymorphism | | Allele classification | Ecotype | Reference |
|---|---|---|---|---|---|---|
| *club-2 (attrs130*[b]*)* | AT5G54440 | SALK_039353* | Intron 14 | Null; seedling[a] lethal | Col-0 | Jaber *et al*, 2010 |
| *trs120-4*[c] | AT5G11040 | SAIL_1285_D07* | Intron 7 | Null; seedling[a] lethal | Col-0 | Thellmann *et al*, 2010 |
| *pp2aa3-1* | AT1G13320 | SALK_014113* | 5'UTR | Null, as evidenced by absence of protein in immunoblot | Col-0 | Zhou *et al*, 2004 |
| *pp2aa3-2* | AT1G13320 | SALK_099550* | Exon 6 | Null | Col-0 | Zhou *et al*, 2004 |

[a]: Seedling - lethal lines were propagated as hemizygotes.
[b]: Note that *club-2* is referred to as *attrs130* in Qi *et al.* (2011).
[c]: This is distinct from the hypomorphic allele later named *trs120-4* by Qi *et al.* (2011).
*: obtained from T-DNA insertion library (Alonso *et al*., 2003).

**Table S2. Primer sequences used for site-directed mutagenesis for TRS120 phosphovariants.** All sequences are shown 5′ to 3′.

| Phospho-variant | Amino acid substitutions | Primer sequences |
|---|---|---|
| SαD | S923D | fwd: 5’-GCC AAG GAA GAT GAT TCT GAC CCA GTA CAA GAT TCT CCA GAG-3’<br>rev: 5’-CTC TGG AGA ATC TTG TAC TGG GTC AGA ATC ATC TTC CTT GGC-3’ |
| SβD | S971D, S973D, S974D, S975D | fwd: 5’-CCC TCC ACC TGG TGA CCC TGA CGA TGA TAG AAA TCC GAG CTT CTC-3’<br>rev: 5’-GAG AAG CTC GGA TTT CTA TCA TCG TCA GGG TCA CCA GGT GGA GGG-3’ |
| SαβγD | S923D, S971D, S973D, S974D, S975D, T1163D, S1165D | fwd: 5’-GCC AAG GAA GAT GAT TCT GAC CCA GTA CAA GAT TCT CCA GAG-3’<br>rev: 5’-CTC TGG AGA ATC TTG TAC TGG GTC AGA ATC ATC TTC CTT GGC-3’<br>fwd: 5’-CCC TCC ACC TGG TGA CCC TGA CGA TGA TAG AAA TCC GAG CTT CTC-3’<br>rev: 5’-GAG AAG CTC GGA TTT CTA TCA TCG TCA GGG TCA CCA GGT GGA GGG-3’<br>fwd: 5’-GTA CTC AGA GCA CGA GCA GGA GAT GCT GAT CCA AAC GAA CCC ATC-3’<br>rev: 5’-GAT GGG TTC GTT TGG ATC AGC ATC TCC TGC TCG TGC TCT GAG TAC-3’ |

**Table S3. Primer sequences used for domestication of PP2A subunits and PTPA for phosphatase activity assay.** All sequences are shown 5′ to 3′.

| Gene | AGI | Primer sequences |
|---|---|---|
| B2 | AT1G17720 | forward_1: GCGCCGTCTCGCTCGTTCGATGAACGGTGGTGACGATGC |
| | | reverse_1: GCGCCGTCTCGTGATCTCCACTTTTATCAAATTCG |
| | | forward_2: GCGCCGTCTCGATCATCTCGCAACTGGTGAC |
| | | reverse_2: GCGCCGTCTCGGCCTCAGATGTTCATGAACC |
| | | forward_3: GCGCCGTCTCGAGGCCCAGGCTATGCGATCT |
| | | reverse_3: GCGCCGTCTCGGGGACGCTCCAAATACGCGG |
| | | forward_4: GCGCCGTCTCGTCCCAAGGAAGTACTGAAGC |
| | | reverse_4: GCGCCGTCTCGCTCACTGCACTGCATAGTACATGTACAAGC |
| FASS | AT5G18580 | forward_1: GCGCCGTCTCGCTCGTTCGATGTATAGCGGATCTAGCGAT |
| | | reverse_1: GCGCCGTCTCGTAGTCTCAAGCTCCATTAGAG |
| | | forward_2: GCGCCGTCTCGACTAAGAGAATCTTGCTCGAG |
| | | reverse_2: GCGCCGTCTCGCATCTCGAACATCTTCTATGC |
| | | forward_3: GCGCCGTCTCGGATGAGATCTGGGACATGGT |
| | | reverse_3: GCGCCGTCTCGCTCAAAGCTACTGAGACTCTTCCTCAGG |
| PUX5/ control | AT4G15410 | forward_1: GCGCCGTCTCGCTCGTTCGATGGCGACGGAGACTAACGAGAAT |
| | | reverse_1: GCGCCGTCTCGGAAGACCGTAGGGAGCAGCG |
| | | forward_2: GCGCCGTCTCGCTTCGATCGAGAGGCGGTGC |
| | | reverse_2: GCGCCGTCTCGCTCACTGCACGAATTTCTGGATGACGACG |
| PP2AC3 | AT2G42500 | forward_1: GCGCCGTCTCGCTCGTTCGATGGGCGCGAATTCTATTCC |
| | | reverse_1: GCGCCGTCTCGGAGTCTCGATAGATGGAGATA |
| | | forward_2: GCGCCGTCTCGACTCTTGACAACATAAGGAATTTTG |
| | | reverse_2: GCGCCGTCTCGCTCAAAGCTACAGGAAATAGTCTGGAGTC |
| PP2AA1 | AT1G25490 | forward: GCGCCGTCTCGCTCGTTCGATGGCTATGGTAGATGAACCG |
| | | reverse: GCGCCGTCTCGCTCAAAGCTAGGATTGTGCTGCTGTGGA |
| PTPA | AT4G08960 | forward: GCGCCGTCTCGCTCGTTCGATGGAACCTCCAAAGGAACAA |
| | | reverse: GCGCCGTCTCGCTCACTGCACCTCTTCCCTGCCAACACC |

**Table S4. piCSi output values from AlphaBridge** (Álvarez-Salmoral et al., 2024) **analysis (related to Fig. S4)**

| | | |
|---|---|---|
| **piCSi:** | **0.83** | |
| **Link** | **Sequence 1** | **Sequence 2** |
| L1 | B2 -21-24 | TRS120-beta peptide -22-26 |
| **piCSi:** | **0.96** | |
| **Link** | **Sequence 1** | **Sequence 2** |
| L1 | PP2AC3 -57-57 | PP2AA1 -489-495 |
| L2 | PP2AC3 -57-57 | PP2AA1 -528-531 |
| L3 | PP2AC3 -76-85 | PP2AA1 -413-413 |
| L4 | PP2AC3 -76-85 | PP2AA1 -450-453 |
| L5 | PP2AC3 -76-85 | PP2AA1 -489-495 |
| L6 | PP2AC3 -76-85 | PP2AA1 -528-531 |
| L7 | PP2AC3 -76-85 | PP2AA1 -535-535 |
| L8 | PP2AC3 -76-85 | PP2AA1 -568-570 |
| L9 | PP2AC3 -113-116 | PP2AA1 -528-531 |
| L10 | PP2AC3 -113-116 | PP2AA1 -568-570 |
| L11 | PP2AC3 -285-287 | PP2AA1 -489-495 |
| L12 | PP2AC3 -285-287 | PP2AA1 -528-531 |
| L13 | PP2AC3 -291-293 | PP2AA1 -450-453 |
| **piCSi:** | **0.9** | |
| **Link** | **Sequence 1** | **Sequence 2** |
| L1 | PP2AC3 -310-313 | PP2AA1 -1-2 |
| L2 | PP2AC3 -310-313 | PP2AA1 -297-297 |
| **piCSi:** | **0.83** | |
| **Link** | **Sequence 1** | **Sequence 2** |
| L1 | PP2AC3 -220-222 | TRS120-beta peptide -25-26 |
| **Link** | **Sequence 1** | **Sequence 2** |
| L1 | B2 -153-158 | PP2AA1 -10-21 |
| L2 | B2 -190-201 | PP2AA1 -10-21 |
| L3 | B2 -190-201 | PP2AA1 -40-43 |
| L4 | B2 -190-201 | PP2AA1 -50-54 |
| **piCSi:** | **0.82** | |
| **Link** | **Sequence 1** | **Sequence 2** |
| L1 | B2 -249-249 | PP2AA1 -1-1 |
| L2 | B2 -259-259 | PP2AA1 -1-1 |

| | | |
|---|---|---|
| **piCSi:** | **0.95** | |
| **Link** | **Sequence 1** | **Sequence 2** |
| L1 | B2 -234-237 | PP2AA1 -179-180 |
| L2 | B2 -234-237 | PP2AA1 -217-219 |
| L3 | B2 -286-287 | PP2AA1 -179-180 |
| L4 | B2 -286-287 | PP2AA1 -217-219 |
| L5 | B2 -286-287 | PP2AA1 -255-258 |
| L6 | B2 -304-307 | PP2AA1 -217-219 |
| L7 | B2 -304-307 | PP2AA1 -255-258 |
| L8 | B2 -304-307 | PP2AA1 -294-297 |
| **piCSi:** | **0.94** | |
| **Link** | **Sequence 1** | **Sequence 2** |
| L1 | B2 -126-127 | PP2AA1 -98-102 |
| L2 | B2 -126-127 | PP2AA1 -140-142 |
| L3 | B2 -253-253 | PP2AA1 -98-102 |
| **piCSi:** | **0.86** | |
| **Link** | **Sequence 1** | **Sequence 2** |
| L1 | B2 -16-24 | PP2AC3 -1-2 |
| L2 | B2 -16-24 | PP2AC3 -124-124 |
| L3 | B2 -16-24 | PP2AC3 -220-220 |
| L4 | B2 -89-90 | PP2AC3 -1-2 |
| L5 | B2 -487-489 | PP2AC3 -1-2 |
| L6 | B2 -500-501 | PP2AC3 -1-2 |
| **piCSi:** | **0.85** | |
| **Link** | **Sequence 1** | **Sequence 2** |
| L1 | B2 -107-108 | PP2AC3 -95-95 |
| L2 | B2 -107-108 | PP2AC3 -133-134 |
| L3 | B2 -107-108 | PP2AC3 -272-273 |
| **piCSi:** | **0.84** | |
| **Link** | **Sequence 1** | **Sequence 2** |
| L1 | B2 -222-222 | PP2AC3 -305-313 |
| L2 | B2 -248-260 | PP2AC3 -305-313 |
| L3 | B2 -307-308 | PP2AC3 -305-313 |
| **piCSi:** | **0.78** | |
| **Link** | **Sequence 1** | **Sequence 2** |
| L1 | A – 153-158 | C – 10-21 |
| L2 | A – 190-201 | C – 10-21 |
| L3 | A – 190-201 | C – 40-43 |
| L4 | A – 190-201 | C – 50-54 |

Swarbreck, D., C. Wilks, P. Lamesch, T. Berardini, M. Garcia-Hernandez, H. Foerster, D. Li, T. Meyer, R. Muller, L. Ploetz, A. Radenbaugh, S. Singh, V. Swing, C. Tissier, P. Zhang, and E.

Huala. 2008. The Arabidopsis Information Resource (TAIR): gene structure and function annotation. *NUCLEIC ACIDS RESEARCH*. 36:D1009-D1014.

Wolny, A., L. Cerrone, A. Vijayan, R. Tofanelli, A.V. Barro, M. Louveaux, C. Wenzl, S. Strauss, D. Wilson-Sánchez, R. Lymbouridou, S.S. Steigleder, C. Pape, A. Bailoni, S. Duran-Nebreda, G.W. Bassel, J.U. Lohmann, M. Tsiantis, F.A. Hamprecht, K. Schneitz, A. Maizel, and A. Kreshuk. 2020. Accurate and versatile 3D segmentation of plant tissues at cellular resolution. *eLife. 9:e57613.*

Wu, T., E. Hu, S. Xu, M. Chen, P. Guo, Z. Dai, T. Feng, L. Zhou, W. Tang, L. Zhan, X. Fu, S. Liu, X. Bo, and G. Yu. 2021. clusterProfiler 4.0: A universal enrichment tool for interpreting omics data. The Innovation. 2:100141.

Zhou, H.-W., C. Nussbaumer, Y. Chao, and A. DeLong. 2004. Disparate roles for the regulatory A subunit isoforms in Arabidopsis protein phosphatase 2A. *The Plant Cell*. 16:709–722.
